# Brd4 and P300 regulate zygotic genome activation through histone acetylation

**DOI:** 10.1101/369231

**Authors:** Shun Hang Chan, Yin Tang, Liyun Miao, Hiba Darwich-Codore, Charles E. Vejnar, Jean-Denis Beaudoin, Damir Musaev, Juan P. Fernandez, Miguel A. Moreno-Mateos, Antonio J. Giraldez

## Abstract

The awakening of the zygote genome, signaling the transition from maternal transcriptional control to zygotic control, is a watershed in embryonic development, but the factors and mechanisms controlling this transition are still poorly understood. By combining CRISPR-Cas9-mediated live imaging of the first transcribed genes (miR-430), chromatin and transcription analysis during zebrafish embryogenesis, we observed that genome activation is gradual and stochastic, and the active state is inherited in daughter cells. We discovered that genome activation is regulated through both translation of maternal mRNAs and the effects of these factors on the chromatin. We show that chemical inhibition of H3K27Ac writer (P300) and reader (Brd4) block genome activation, while induction of a histone acetylation prematurely activates transcription, and restore genome activation in embryos where translation of maternal mRNAs is impaired, demonstrating that they are limiting factors for the activation of the genome. In contrast to current models, we do not observe triggering of genome activation by a reduction of the nuclear-cytoplasmic (N/C) ratio or slower cell division. We conclude that genome activation is controlled by a time-dependent mechanism involving the translation of maternal mRNAs and the regulation of histone acetylation through P300 and Brd4. This mechanism is critical to initiating zygotic development and developmental reprogramming.

---

Upon fertilization, the genome of metazoan embryos remains transcriptionally silent until the maternal-to-zygotic transition (MZT). Maternally-deposited proteins and RNAs regulate zygotic genome activation (ZGA), and failure of transcription activation results in developmental arrest^1-4^. The timing and the number of divisions that precede genome activation is highly reproducible within each species, suggesting a robust temporal regulation, yet the mechanisms that control when and how the genome becomes activated remain poorly understood.

Studies in different species have proposed three models to explain how the zygotic genome becomes activated: slowing of the cell cycle (slow cycle model)^5-7^, the nuclear-cytoplasmic (N/C) ratio (N/C ratio model)^8-10^ or an elapsedtime-dependent mechanism(developmental timer model)^11-13^. The slow cycle model posits that during early development, rapid cell divisions have been postulated to inhibit productive transcription, due to competition between the replication and transcription machineries^7,11,14^. Indeed, global transcriptional activation coincides with the slowdown of the cell cycle^15-19^. Early transcribed genes are the first to be replicated in zebrafish^6^, and increasing the replication time can increase transcription in *Xenopus* and zebrafish^5,20^. However, it is unclear whether the slowing of the cell cycle is the main trigger for genome activation.

The N/C ratio model, originated in *Xenopus,* proposes that a predetermined N/C ratio could establish a permissive condition for genome activation^2,9,10^. Consistent with this model, manipulation of ploidy in other species alters the timing of detection for transcription in some genes^15,16,21,22^. However, these perturbations do not affect all genes^16,21,23^, and the changes in the amount of template DNA could result in lower sensitivity for transcribed genes in haploid embryos, explaining the delayed transcription observed in haploid embryos^12^. The exponential increase in DNA content is thought to titrate maternally-deposited transcriptional repressors^8-10^, namely histones, which are thought to compete with transcription factor binding and block access of the transcriptional machinery to DNA^24-28^. *In vitro* studies have demonstrated a repressive effect of histones on transcription using *Xenopus* egg extract^8-10^, yet their effect in vivo for regulating genome activation is modest^8,26,29^. Indeed, it is clear that the N/C ratio model alone cannot explain how the genome becomes activated, and haploid *Drosophila* embryos activate the expression of many zygotic genes in a time-dependent manner^15,16,23^.

According to the developmental timer model, the translation of maternally deposited mRNAs that regulate transcription or the epigenetic marks controls the timing of genome activation. During MZT the genome acquires bivalent histone marks H3k4me3 and H3k27me3^30-33^, remodels the DNA methylation pattern^34-37^ and acquires new nucleosomes in the promoter of zygotic genes^38,39^. Maternally-deposited transcription factors such as Zelda in *Drosophila*^40-43^ and Pou5f3, Sox19b, Nanog in zebrafish^44,45^, are highly translated and required for activating the first zygotically expressed genes. Similarly, translation of TBP RNA contributes to the transcriptional capacity during genome activation in *Xenopus* and zebrafish^46-48^. While the changes at the level of transcription and the chromatin are well characterized, which factors act as developmental timers and how they initiate genome activation during embryogenesis remain unknown. Here, we use imaging and sequencing analysis in zebrafish to interrogate the molecular nature of the timer and find that histone acetylation through p300 and Brd4 regulate activation of the genome after fertilization.

## Results

### Genome activation during MZT initiates stochastically in the miR-430 locus

The maternal-to-zygotic transition (MZT) involves a major remodeling of the transcriptome after fertilization^14,44^. To define the set of zygotic genes activated during this transition we conducted pull down of nascent transcripts with 5-ethynyl uridine (EU) incorporation and RNA-seq (Click-iT-seq)^14,44^, and identified 2669 zygotically transcribed genes by 4hpf (Extended Data Fig. 1 and Methods). This represents a major switch in transcriptional competency of the genome. Yet, it remains unclear how transcription begins, whether it occurs simultaneously or gradually across the genome, and whether all the cells activate the genome simultaneously. To address these questions, we analyzed global transcription at a single-cell resolution during MZT. We assayed for poised RNA polymerase II (RNAPol II phospho Ser5; Pol II p-Ser5) (Fig. 1b,e) and for RNA synthesis by injecting EU at the one-cell stage, which incorporates into nascent RNA and is detected by Click-It chemistry through imaging (Fig. 1a,d). We first observed poised RNA pol II and nascent transcription in two foci at the 64-cell stage, 2 hours post-fertilization (hpf) (Fig. 1b, d, e). Transcription increased gradually over time, and became broadly distributed within the nucleus by sphere stage (4 hpf) (Fig. 1d, e). As a control, embryos treated with RNAP II inhibitors triptolide^49^ or α-amanitin^50,51^ lacked EU labeling of nascent transcripts (Fig. 1d). Pol II p-Ser5 signal colocalized with foci of active transcription and was limited to specific stages of the cell cycle during late interphase and early prophase (Extended Data Fig. 2b), indicating that recruitment and poising of Pol II is regulated during the cell cycle.

**Figure 1.**
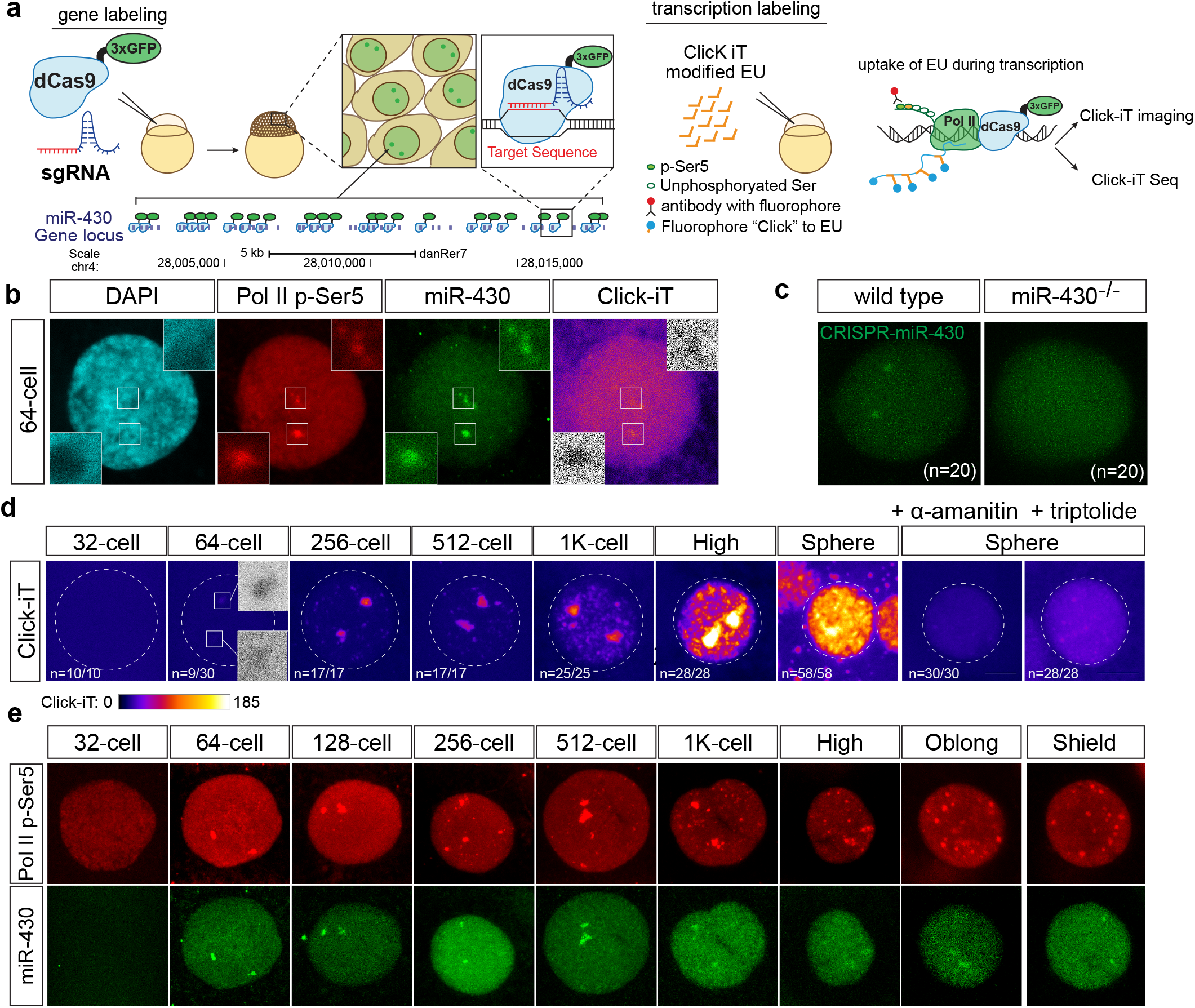
Zygotic genome activation occurs gradually and begins with the miR-430 locus. **(a)** Schematic illustrating strategies used to visualize the miR-430 locus using CRISPR-dCas9-3xGFP (dCas9) (left) and global transcription using Click-iT chemistry (right). **(b)** Representative single nucleus confocal images labeled with DAPI, phosphorylated Ser5 RNA pol II (Pol II p-Ser5), miR-430 loci, and Click-iT (transcription). Note the co-localization of the earliest detectable Pol II p-Ser5, Click-iT signal with CRISPR-dCas9 labeled miR-430 locus at 64-cell stage (~2hpf). Insets represent single plane images of individual foci. **(c)** CRISPR-dCas9 labeling reveals two foci signals in wild-type embryos but not in miR-430^−/-^ mutants that lack the miR-430 locus, demonstrating the signal specificity towards the miR-430 endogenous locus. (n=number of analyzed nuclei across three embryos). **(d)** Time-course imaging analysis of Click-iT labeled zygotic transcription from 32-cell (~1.75hpf) to sphere stage (~4hpf). The earliest Click-iT signal was detected in two foci at the 64-cell stage. Representative nucleus from embryos treated with transcription inhibitors (α-amanitin and triptolide) are shown as negative control. Scale bar represents 5m. Nucleus is outlined by a dotted line. DAPI staining is used for staging nucleus at comparable stages within the cell cycle for subsequent imaging analysis conducted in this study (Extended data Fig. 2b). **(e)** Time-course analysis of single nuclei labeled for Pol II p-Ser5 and dCas9 targeting the miR-430 locus from 32-cells to shield stage. Note that the first detectable Pol II p-Ser5 foci signal co-localizes with miR-430 loci at 64-cell stage. As transcription is globally activated, multiple Pol II p-Ser5 foci appear across the nucleus.

To identify the first zygotically-transcribed genes, we combined whole transcriptome analysis with CRISPR-dCas9-GFP-mediated labeling of endogenous loci (Fig. 1a-c,e and Extended Data Fig. 2). RNA-seq pointed to miR-430 as a potential candidate for the first transcribed locus^14,52^ (Extended Data Fig. 1b,c). To test this, we co-injected dCas9-3xGFP with 2gRNAs targeting 20 sites on the endogenous miR-430 locus. We observed that miR-430 loci co-localize with the earliest detected transcription and pSer5PolII signal in 64-cell stage embryos (Fig. 1b, e; Extended Data Fig. 2d). This signal is highly specific to miR-430, as it is not detected in a miR-430 deletion mutant (Fig. 1c). These results indicate that the genome undergoes a gradual activation of transcription during MZT, in which miR-430 is the earliest active locus detected in the genome.

During early development, rapid cell cycles are synchronous^53^. To determine whether genome activation occurs simultaneously throughout the embryo, we developed a method to visualize transcription *in vivo* using a molecular beacon (MB^miR430^) complementary to the primary miR-430 transcript (Fig. 2a). The 5’ and 3’ ends of the stem are modified with a Lissamine fluorescent tag and a fluorescence quencher (Dabcyl), which reduces the fluorescent background when the probe is not bound to its target. This molecular beacon detects miR-430 transcription specifically, as the signal is lost in miR-430 deletion mutants^54^ and in embryos treated with the Pol II inhibitor α-amanitin (Fig. 2b;Extended Data Fig. 3a). Injection of MB^miR430^ into one-cell stage embryos, followed by timelapse analysis, revealed that transcriptional competency is briefly achieved towards the end of cycle 6 (by the 64-cell stage) before the cell cycle slows down (Fig. 2c-f), with an increase in the intensity and duration of transcription over each cell cycle (Fig. 2e, f). Despite the synchronous cell cycle across the early embryo, lineage analysis of miR-430 transcription revealed that transcription competency is achieved in a stochastic pattern across the cells (Fig. 2g; Extended Data Fig. 3b). Most cells activated both alleles simultaneously (>80% of the cells), and once competency is achieved, the active state is maintained in the daughter cells (Fig. 2g, Extended Data Fig. 3b). These results suggest that transcriptional competency is first achieved in the miR-430 locus, in a stochastic and inherited pattern during development.

**Figure 2.**
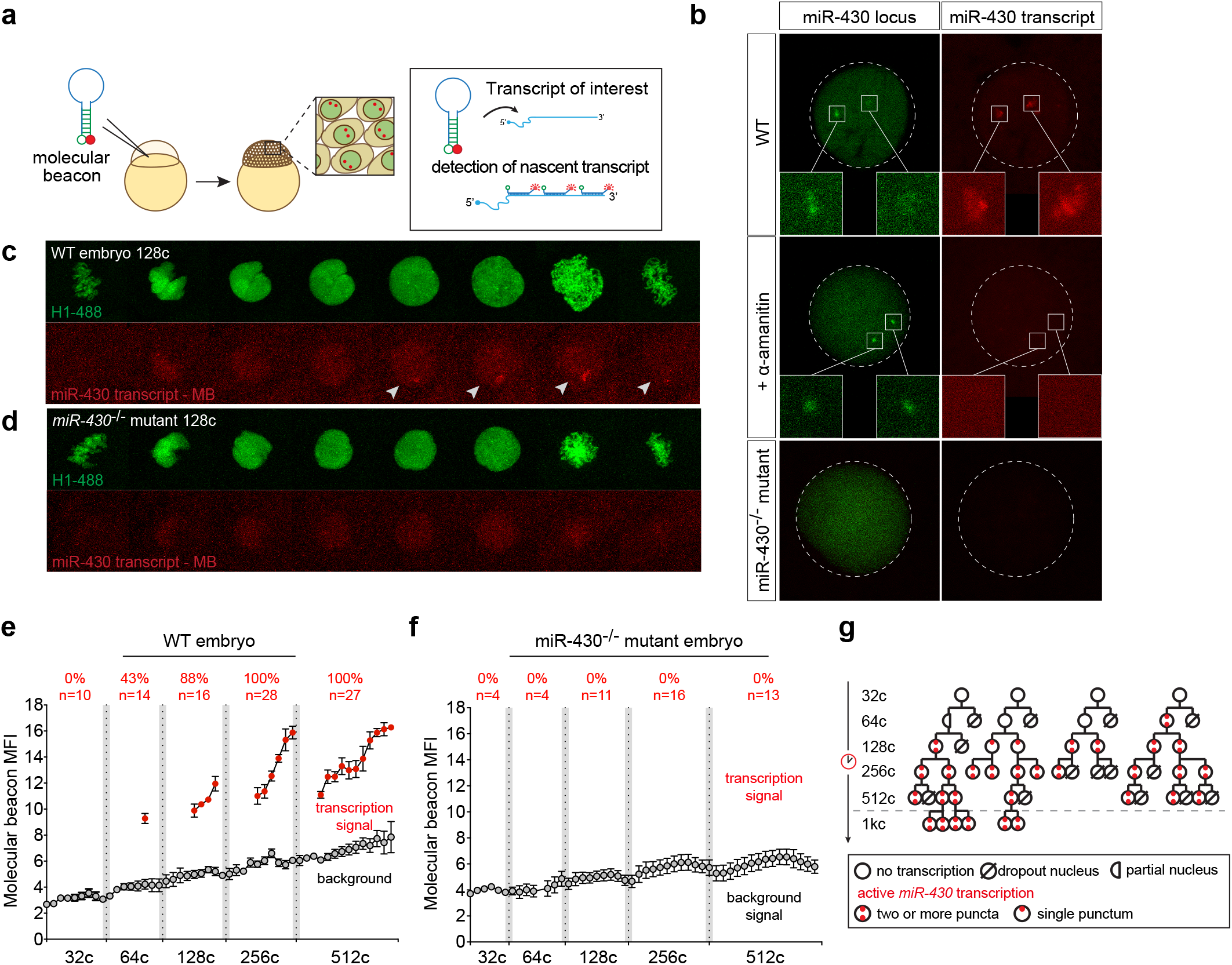
Live imaging of miR-430 transcription reveals that transcription competency is acquired in a stochastic and heritable manner. **(a)** Schematic illustrating *in vivo* labeling of nascent transcript of miR-430 in zebrafish embryos for live imaging using a molecular beacon (MB^miR430^). **(b)** Confocal imaging of wild-type, α-amanitin-treated, and miR-430^−/-^ mutant embryos labeled with MB^miR430^ (red) and dCas9-miR-430 (green). Note the absence of miR-430 transcription in miR-430^−/-^ deletion mutants and after Pol II inhibition. **(c, d)** miR-430 transcription *in vivo* visualized by MB^miR430^ during 128-cell stage cell cycle in wild type (c) or miR-430^−/-^ mutant (d). Chromatin is labeled with Alexa Fluor^®^ 488 histone H1. MB^miR430^ signals are highlighted by white arrows in the confocal images. miR-430 transcription is only detected in late interphase and early prophase. Note the absence of MB^miR430^ signal in miR-430^−/-^ mutants. **(e)** MB^miR430^ mean fluorescence intensity (MFI) quantified over each cell cycle in wild-type embryos and **(f)** miR-430^−/-^ mutant embryos. Red dots represent the time points when MB^miR430^ signal is detected; grey dots represent the nuclear background MB^miR430^ signal. (n=number of nuclei imaged; percentage represents the proportion of analyzed nuclei that display miR-430 transcription) **(g)** Representative cell-lineage tracing for miR-430 transcription. Transcription competency is acquired in a stochastic manner at 64c, and once gained is maintained in the daughter cells after cell division.

### Transcriptional competency depends on a developmental timer independent of cell division

Contrary to the proposed slow cycle model of embryonic genome activation, our analysis of transcription in live embryos demonstrates that transcriptional activation of miR-430 initiates before the cell cycle slows down (Fig. 2e). To further analyze the expression of other genes in the genome and distinguish between the aforementioned models, we uncoupled developmental time and cell division by blocking DNA replication (Fig. 3a) and analyzed global transcription using Click-iT-seq (Extended Data Fig. 1a). *Chk1* inhibits *Drf1* and blocks the formation of the origin of replication which in turn slows down cell division during the mid-blastula transition (MBT)^55^. Premature *chk1* expression through mRNA injection at the 1-cell stage stops cell division in *Xenopus*^55^ and in zebrafish (Fig. 3b, c), arresting the embryos between 4-and 16-cell stages throughout the first 6 hours of development. Analysis of chk-1 injected embryos (chk1^OE^) at 2 hpf revealed that slowing down the cell cycle can increase miR-430 transcription early on (Extended Data Fig. 4e), consistent with the regulation of Pol II during the cell cycle^56^ (Extended Data Fig. 2b). However, it is not sufficient to cause widespread activation of the genome at this stage as assayed by Click-iT labeling of transcription (Extended Data Fig. 4e). These results suggest that slowdown of the cell cycle during MBT can contribute to the transcriptional output but is not sufficient to trigger premature genome activation.

**Figure 3.**
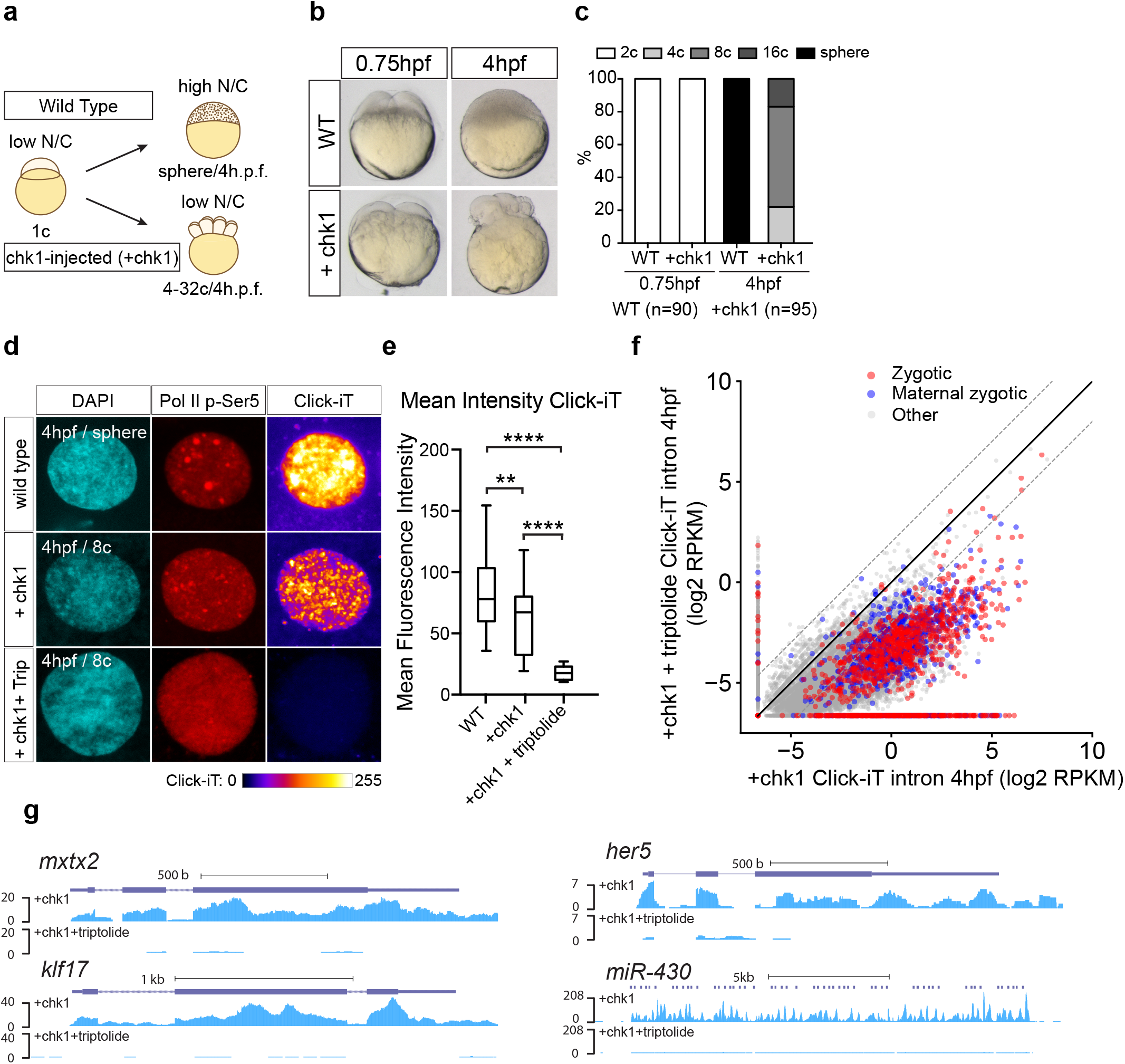
Genome activation occurs over time independent of the N/C ratio. **(a)** Schematic illustrating the strategy to induce cell cycle arrest by expressing chk1 in zebrafish embryos **(b)** Transmitted light microscopy picture of wild type (WT) and chk1-injected embryos (+chk1) at different times (hours post fertilization, hpf). Note how *chk1* expression inhibits cell divisions in zebrafish embryos at 4hpf. **(c)** Bar plot quantifying the number of cells in wild type (WT) and chk1 (n=number of embryos analyzed) **(d)** Single nucleus confocal image of DAPI, Pol II p-Ser5 and transcription (Click-iT) in different conditions as indicated at 4hpf (+Trip, incubated with triptolide). Click-iT signal intensity is presented in a heatmap color scale**. (e)** Box and whisker plots showing the mean fluorescence intensity for Click-iT signal in the conditions shown in **(d)**. (\*\*\*\**P*<0.0001; ** *P*=0.0058, two-sample t-test: WT (n=129); +chk1 (n=21); +chk1+triptolide (n=15)) **(f)** Biplot comparing intron expression levels of genes in chk1-injected embryos with and without triptolide treatment reveals genome activation occurs in chk1-injected embryos at 4hpf. The solid black line represents the diagonal and the dashed lines represent 4-fold change. **(g)** Genome tracks representing normalized Click-iT-seq signal for examples of zygotic genes activated in chk1-injected embryos with (+chk1+triptolide) and without triptolide (+chk1).

Chk1^OE^ embryos also allowed us to address the titration of maternal repressors by successive cell divisions, testing the N/C ratio model of embryonic genome activation. Quantification of the DNA and histone levels demonstrated that chk1^OE^ embryos stall cell division, and maintain a low N/C ratio over time, with DNA template and histone levels equivalent to 16-32-cell stages of wild-type embryos (Extended Data Fig. 4a, b). Despite their low N/C ratio, chk1^OE^ embryos activated their genomes later on at 4 hpf compared to triptolide treated embryos as shown by EU labeling of transcription and Click-iT-seq (Fig. 3d-g; Extended Data Fig. 4c, d). These results suggest that titration of maternal histones by the increase in DNA content is not required for genome activation. We analyzed zygotic transcription using Click iT-seq of nascent RNAs to quantify both exonic and intronic sequences, where the latter provides better signal to noise ratio as most maternally deposited mRNAs are spliced and the introns are degraded early on^44^. This increases the sensitivity of Click-iT-seq, especially in embryos with low number of cells such as chk1^OE^ embryos. Click-iT-seq revealed that chk1^OE^ embryos activate 67.8% of the zygotic genes compared to triptolide treated chk1^OE^ embryos (1218 zygotic and 591 Maternal zygotic genes) (Fig. 3f and g). Transcription levels in chk1^OE^ embryos are moderately lower than wild type as assayed by single nucleus Click-iT labeling of transcription (Fig. 3e). Our results are in direct contrast with the N/C model, which predicts that genome activation cannot occur without DNA replication to increase the N/C ratio. Instead, our work demonstrates the developmental timer model to initiate zygotic transcription.

### Translation of maternally-provided mRNAs controls ZGA

We hypothesized that the timer regulating transcriptional competency depends on the translation of maternal mRNAs. To test this model, we blocked translation initiation and elongation with Pateamine A and Cycloheximide^57-59^ at the 8-or 32-cell stages, which arrests the cell cycle at the 16-and 64-cell stages, respectively (Fig. 4a). Click-iT seq of zygotic transcription revealed global transcriptional activation by 4 hours when translation was inhibited by the 64-cell stage when compared to α-amanitin treated embryos (2213 genes upregulated ≥4-fold, see Methods, Fig. 4b, c; Extended Data Fig. 5). Examples of these include *mxtx2*, *klf17*, *her5* and *aplnrb* (Fig. 4d). On the contrary, blocking translation 30 minutes, by the 16-cell stage, resulted in a notable reduction in transcriptional competency (256 genes upregulated ≥4-fold, see Methods, Fig. 4b, c; Extended Data Fig. 5). Based on these results, we conclude that the maternal factors translated by the 64-cell stage (2 hpf) are sufficient to trigger global genome activation for 82.9% of the zygotic transcripts. Despite the constant N/C ratio over that time, global activation is only observed at 4 hpf, suggesting that the maternal factors translated by the 64-cell stage still require additional time to induce transcriptional competency, possibly by regulating the chromatin during that time.

**Figure 4.**
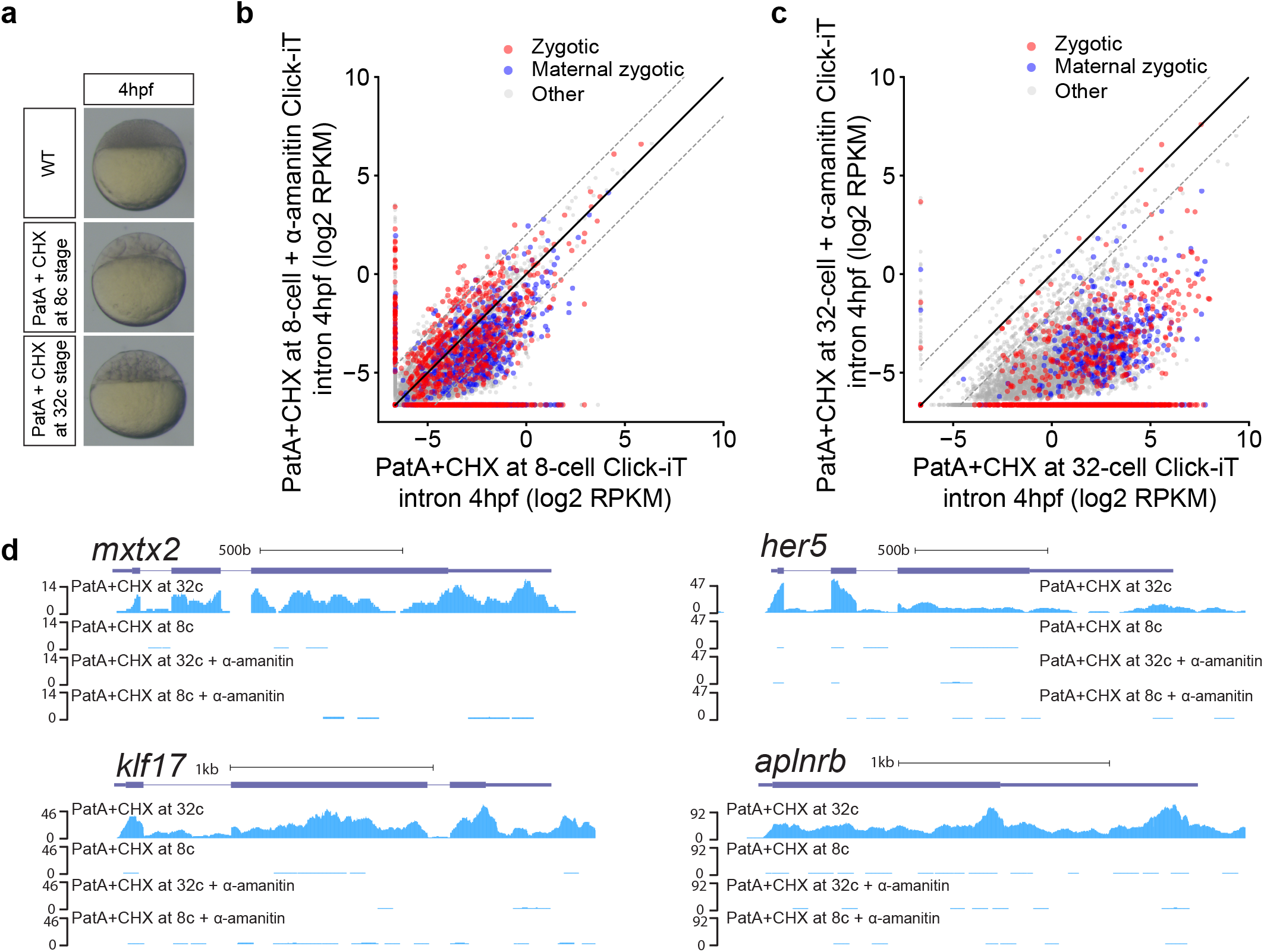
Transcriptional competency of the zygotic genome depends on the translation of maternal mRNAs. **(a)** Embryos treated with pateamine A (PatA) and cycloheximide (CHX) at 8-(1.25hpf) and 32-cell stage (1.75hpf) arrests zebrafish embryos at 16-and 64-cell stage respectively. **(b, c)** Biplot of Click-iT-seq RNA levels at 4hpf comparing the level of transcription using intron signal. Embryos were treated with PatA + CHX at 8-cell stage (b) or at 32-cell stage (c), with and without α-amanitin. Note the limited level of transcription when translation is blocked at 8-cell stage, compared to the increase in transcription when translation is blocked 30 minutes later at 32-cell stage. Dashed lines represent 4-fold change**. (d)** Genome tracks representing normalized Click-iT-seq signal in the embryos described in (b) and (c). Click-iT-seq was normalized by the total number of mitochondrial RNAs, as an internal control (Methods).

### Acquisition of H3K27Ac correlates with transcriptional activation

To identify temporal regulators of genome activation, we analyzed the chromatin marks labeling the first active genes and interrogated their function using chemical inhibitors. Two lines of evidence indicate that genome activation coincides with the acquisition of H3K27Ac and requires reading of this mark by bromodomain-containing proteins. First, we analyzed the first active gene (miR-430) for histone modifications H3K27Ac, H3K4me1, H3K4me3, at dome stage (4.3 hpf), using public datasets^60^. miR-430 was the top labeled locus with each of these marks (Fig. 5a, Extended Data Fig. 6a-c). Consistent with these results, imaging analysis revealed that the earliest H3K27Ac and H3K4me3 signals co-localized with the miR-430 locus (Fig. 5c, Extended Data Fig. 6g). At later stages, among the different histone marks, H3K27Ac showed the best correlation with transcription as assayed by Click-iT-seq (r=0.53, Spearman correlation, P = 3.9×10^−50^, rank correlation independence test) (Fig. 5b; Extended Data Fig. 6d-f). Second, we analyzed the effect of different small molecule inhibitors targeting readers or writers for H3K4me3 and H3K27Ac (Extended Data Fig. 6h). Among these drugs, treatment with JQ1 (an inhibitor of BET bromodomains BRD2-4), or SGC-CBP (an inhibitor of histone acetyltransferase P300 and CBP) resulted in a significant reduction of zygotic transcription (Fig. 5d-g, Extended Data Fig. 7), which in turn blocked gastrulation (Fig. 5d). For example, intron analysis revealed that 84% of zygotically transcribed genes were reduced in expression ≥4-fold by the bromodomain inhibitor JQ1 when compared to wild type embryos (Fig. 5f). This effect was also apparent for individual transcripts corresponding to both housekeeping and developmental genes (Fig. 5g; Extended Data Fig. 7g). These results suggest that the targets of JQ1 and SGC-CBP (BRD4 and P300/CBP) are required to initiate zygotic genome activation.

**Figure 5.**
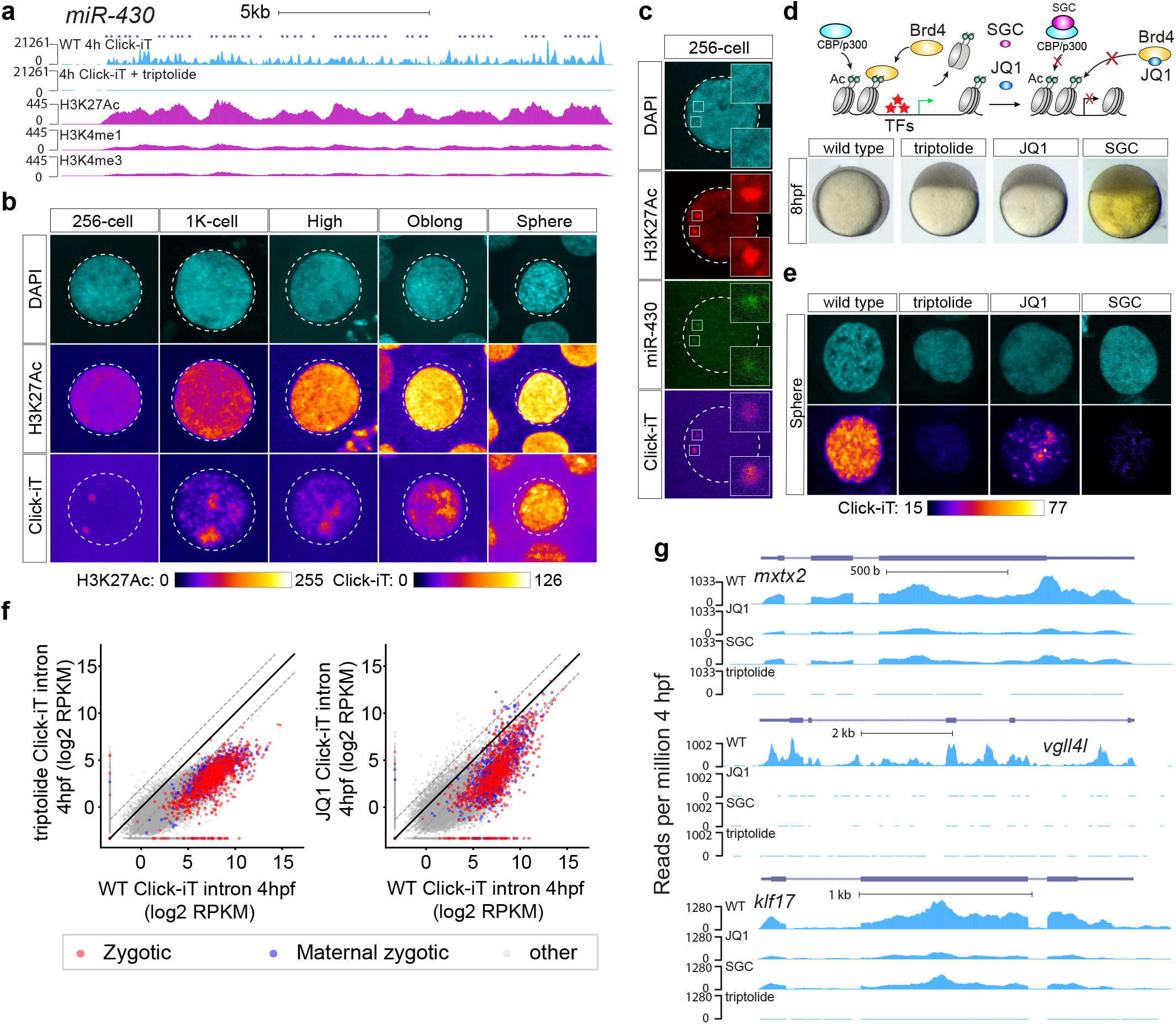
H3K27Ac correlates with transcriptional activation during ZGA. **(a)** Genome tracks representing normalized Click-iT-seq signal and histone mark level at the miR-430 locus. **(b)** Time resolved single nucleus confocal imaging analysis of H3K27Ac from 256-cell to sphere stage reveals a positive correlation with the level of Click-iT signal. Both H3K27Ac and Click-iT signal intensity are presented in a heatmap color scale. **(c)** Single plane confocal image labeled for DAPI, H3K27Ac, dCas9-miR-430 and Click-iT. Note the co-localization of H3K27Ac with Click-iT labeled transcription activity at the miR-430 locus. **(d)** Schematic illustrating the selective pharmacologic inhibition of the acetyl-lysine-binding pocket of the BET bromodomain proteins (BRD2-4) and CBP/P300 by JQ1 and SGC respectively (top). Embryos treated with JQ1 and SGC both arrest before gastrulation similar to those treated with triptolide, consistent with a loss of zygotic transcription (bottom) (Ac = H3K27 acetylation, TFs = transcription factors). **(e)** Click-iT imaging analysis in wild-type, triptolide, JQ1 and SGC treated embryos reveals a significant reduction in transcription by the treatment of JQ1 and SGC. Click-iT signal intensity is presented in a heatmap color scale. **(f)** Biplot comparing intron expression levels of genes measured by Click-iT-seq in triptolide (left) and JQ1 (right) treated embryos with wild-type embryos at 4hpf. Note the strong reduction in genome-wide transcription with JQ1 and SGC (Extended Data Figure7). Dashed lines represent 4-fold change. **(g)** Genome tracks representing normalized Click-iT-seq signal measured at 4hpf in wild-type, triptolide, JQ1 and SGC treated embryos for examples of zygotic genes.

Next, we analyzed whether the reader (BRD4) and writer (P300) of histone acetylation can regulate the timing of genome activation. Injection of P300 and BRD4 proteins at the one-cell stage led to premature activation of the zygotic genome; this was manifested by an increase in H3K27Ac that coincided with Click-IT signal, revealing premature miR-430 expression at the 32-cell stage (Fig. 6a). Click-iT analysis of zygotically transcribed mRNAs revealed an increase in transcriptional output and H3K27Ac, by 1K-cell stages (Fig. 6b, c; Extended Data Fig. 8). In support of this, RNA-seq analysis revealed >2 fold increase in zygotic transcription in p300+BRD4 expressing embryos compared to wild type (Fig. 6d, e, f), with 729 genes upregulated ≥4-fold in P300+BRD4 injected embryos.

**Figure 6.**
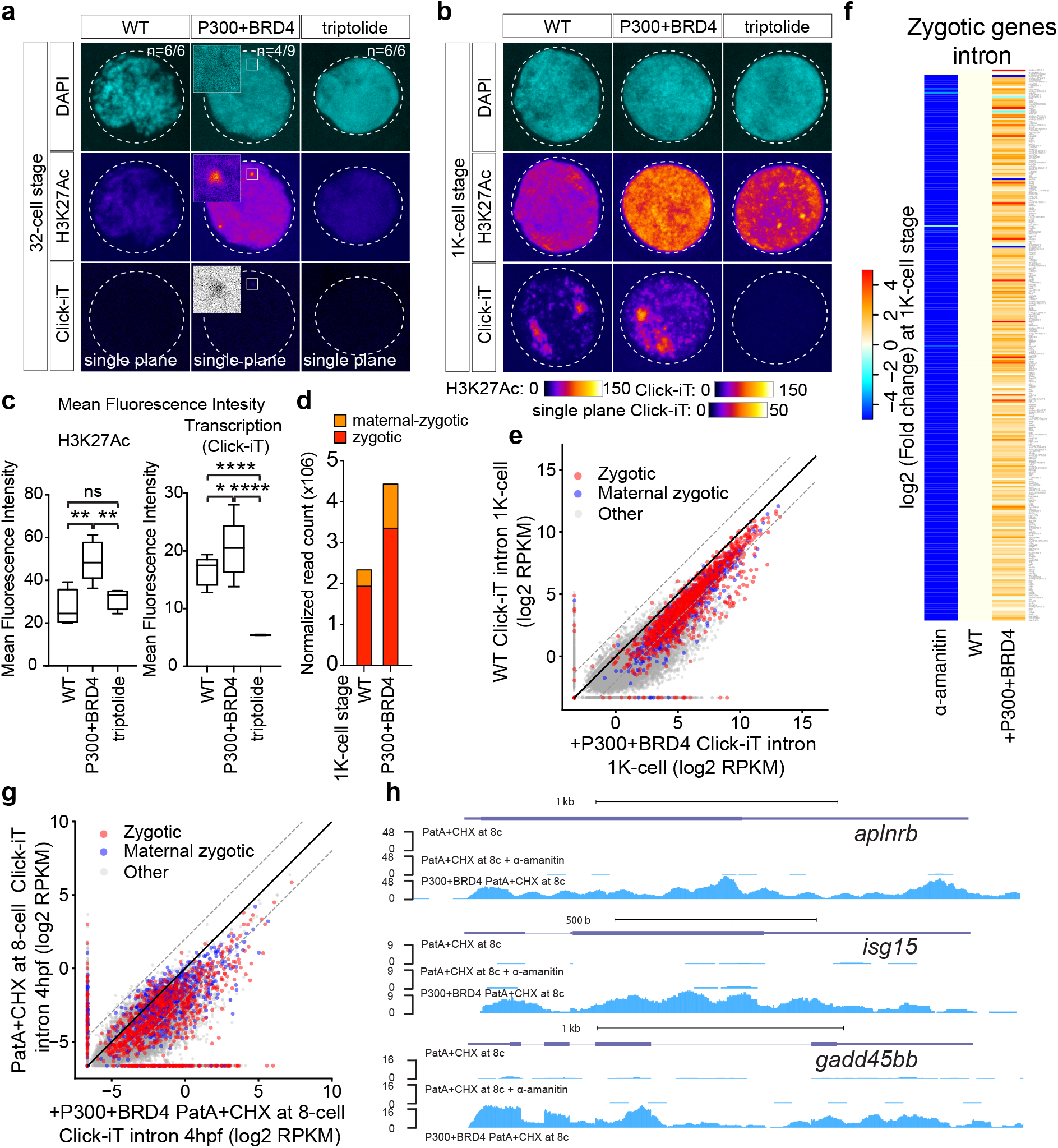
P300 and BRD4 are limiting factors regulating zygotic genome activation. **(a, b)** Single nucleus imaging analysis of H3K27Ac and Click-iT signal comparing embryos with and without early expression of P300 and BRD4 at 32-cell **(a)** and 1K-cell stage **(b)**. Note the increased acetylation and premature transcription by early expression of P300 and BRD4 proteins (WT: wild-type embryos; P300+BRD4: embryos with early expression of P300 and BRD4 protein; triptolide: embryos treated with triptolide). Both H3K27Ac and Click-iT signal intensity are presented in a heatmap color scale. Inset highlights the single plane confocal image of region with Click-iT signal co-localizing with high H3K27Ac signal. (n=number of incidence among the total number of nuclei imaged) **(c)** Box and whisker plots quantifying the mean fluorescence intensity for H3K27Ac and Click-iT signal in the conditions described in **(b)** (H3K27Ac signal: P300+BRD4 > WT *P*=0.0068; P300+BRD4 > triptolide *P*=0.0013; WT~Trip *P*=0.2482. Click-iT signal: P300+BRD4 > WT *P*=0.0068; P300+BRD4 > triptolide *P*=0.0013; WT~Trip *P*=0.2482). **(d)** Stacked bar plots comparing the total number of read count (normalized by total read count of mitochondrial protein-coding genes) of zygotic and maternal zygotic genes in wild-type embryos (WT) and embryos with early expression of P300 and BRD4 protein (P300+BRD4) at 1K-cell stage. **(e)** Biplot comparing intron expression levels measured with Click-iT-seq in wild-type embryos and embryos with early expression of P300 and BRD4 at 1K-cell stage. Note the upregulation of genes expressed in embryos with early expression of P300 and BRD4. Dashed lines represent 4-fold change. **(f)** Heatmap comparing intron expression levels of published zygotic genes (Methods) in α-amanitin treated embryos (α-amanitin) and embryos expressing P300 and BRD4 (+P300+BRD4) with wild-type embryos (WT). Genes in the heatmaps are ranked by transcription level from low to high (top to bottom) based on the difference in intron expression levels (intron RPKM) between wild-type embryos and α-amanitin treated embryos. Genes without intron are not included in the heatmaps. **(g)** Biplot comparing intron expression levels measured by Click-iT-seq for embryos treated with PatA + CHX at 8-cell stage injected with and without P300 and BRD4. Note that early expression of P300 and BRD4 protein restores transcription competency in these embryos. **(h)** Genome tracks representing normalized Click-iT-seq signal in the conditions described in (g). Normalized Click-iT-seq signal for embryos treated with PatA + CHX at 8-cell stage in the presence of α-amanitin is shown as control.

The modulation of translation during development demonstrated that maternal factors are limiting early in development, hence preventing transcriptional competency. To determine whether P300 and BRD4 are sufficient to regulate transcription when translation of maternal mRNAs is blocked, we introduced P300+BRD4 into PatA+CHX-treated embryos by the 16-cell stage. Quantification of zygotic transcription using Click-iT revealed an increase in zygotic transcripts compared to wild type embryos, with 623 genes activated ≥4-fold in the presence of P300 and BRD4 (Fig. 6g, h, Extended data Fig. 9). Together, our results suggest that acetylation components P300 and Brd4 are limiting factors that regulate the timing and transcriptional output during zygotic genome activation.

## Discussion

Our results provide two major insights on the mechanisms that activate the zygotic genome after fertilization. First, activation of the genome largely depends on the translation of maternal mRNAs over developmental time, independent of the number of cell divisions or the N/C ratio. The N/C ratio has long been proposed in different model organisms (*Xenopus*, zebrafish and *Drosophila*) as a mechanism that regulates the mid-blastula transition (MBT)^2,17,26,29,61,62^. This transition includes changes in cell-cycle length, checkpoint activation, and transcription of zygotic genes^2,17,26,29,61,62^. Indeed, it is clear that N/C ratio controls specific aspects of MBT, as the slowdown of the cell cycle depends on the titration of the replication factors by the DNA and the inhibition of Drf1 by Chk1^55^. However, our results reveal that transcriptional activation is not triggered by the titration of cytoplasmic repressors by replicating DNA.

Instead, we propose that the embryo measures time through the translation of maternal mRNAs and the effects of these products on the chromatin, which depend on elapsed time. Consistent with this model, haploid embryos in *Drosophila* activate a large set of genes independent of the N/C ratio^15,22^, and different chromatin regions acquire accessibility independent on the N/C ratio^23^, while other regions appear to depend on an N/C timer^23^. Histones have been shown to repress transcription in vitro^29^. It has been proposed that maternally loaded histones, which are in great excess, compete with specific transcription factors and repress premature genome activation^26^. It is possible that the increase in histone acetylation observed during MZT relieves the repressive activity of histones. Genetic modulation of histone levels will be needed to further address their role in transcriptional repression.

Rapid cell division has also been proposed to regulate genome activation, and miR-430 has been identified as an early replicating locus in the zebrafish genome^6^. We observe that Pol II is regulated during the cell cycle (Extended data Fig. 2). This is consistent with the increase in transcription output with the lengthening of the cell cycle^5,20^, suggesting that the length of the cell cycle contributes to the global transcriptional output during MBT. However, slowing the cell cycle does not lead to a significantly premature activation of the genome. The fact that miR-430 transcription begins before the slowdown of the cell cycle suggests that lengthening of the cell cycle is not the main determinant triggering genome activation (Fig. 2c-f), but could contribute to transition from stochastic to global transcription^63^. Indeed, early gene expression is also observed in *Drosophila* before the cell cycle slows down^64^. It is possible that a small number of reductive divisions are needed to allow genome activation, yet we observed that beyond 32 cells, the timing of activation is independent of cell division. Our results provide direct evidence for the requirement of elapsed time for genome-wide transcription, independent of the cell cycle length or the N/C ratio, as the main mechanism controlling genome activation.

Second, we demonstrate that a histone acetylation writer (P300) and reader (BRD4) modulate genome activation during MZT, as chemical inhibition and premature expression of P300 and BRD4 can change the timing of genome activation. Consistent with these results, it has been shown that histone acetylation of H4K8Ac, H3K18Ac, and H3K27Ac coincide with the activation of the genome in *Drosophila*^65,66^. However, since ectopic H3K27Ac only induces premature activation of zygotic genes consistent with physiological chronology, additional activators and remodeling events must be required; the acquisition of H3K27Ac alone is not sufficient for transcriptional activation when other activators remain at a limiting protein threshold. Previous studies in zebrafish have shown that sequence-specific transcription factors (TFs) Nanog, SoxB1 and Pou5f1 are required for initiating ~75% of the first major wave of ZGA^44,45^. Other studies, in mouse embryos, have demonstrated that other regulators, such as the DUX transcription factor and the chromatin remodeler BRG1, are required for establishing chromatin structure and transcription competence during ZGA^67-71^. However, overexpression of only Nanog, SoxB1, Pou5f1, and Brg1 did not yield premature activation similar to that of p300 and BRD4 (Extended Data Fig. 8a), suggesting a different temporal requirement among these activators. At the moment the temporal relationship between histone acetylation and transcription factor binding is not resolved^65,66^. Answering this question and determining the role of chromatin accessibility will require performing loss of function studies for different TF activators during MZT, combined with high resolution imaging or ChiP methods amenable for low input. The observed premature activation of miR-430 at the 32-cell stage in the context of p300 and BRD4 early expression (Fig. 6a) suggests that the miR-430 locus is properly primed during early stages, but additional factors are likely limiting at these stages to allow global activation of the genome. Studies in ES cells have shown that Nanog interacts with p300 and BRD4, and there is a tight correlation of binding and recruitment between these factors^72,73^. Furthermore, P300 and BRD4 can restore transcriptional competency in embryos with reduced translation. Hence, we propose P300 and BRD4 are limiting factors for the activation and serve as downstream effectors of transcription activation during ZGA.

Multiple factors involved in genome activation also function during cellular reprogramming and stem cell maintenance^12,74^. For example, the homologue of miR-430 in mouse, miR-295/miR-302, facilitates reprogramming of differentiated cells. miR-295/miR-302 expression is controlled by a super enhancer in ES cells, and miR-430 is the strongest labeled gene with H3K27Ac and H3K4me1, which typically mark super enhancers. These parallels are interesting, since one of the first events the embryo must accomplish is the reprogramming of the differentiated sperm and oocyte nuclei into a transient totipotent state receptive to various differentiation programs. Given the common regulatory programs between miR-430 and miR-302/295, including Brd4 and p300, Nanog and Oct4, it will be interesting in the future to determine whether other factors controlling super enhancer function also regulate genome activation in vertebrates.

In summary, the results presented here open a new dimension into understanding the maternal factors and mechanisms that enable activation of the genome – by regulating histone acetylation. Analyzing transcription of the first active genes in live embryos where most of the genome is silent can help us determine how chromatin active regions are first organized at the single cell level, and how different factors bind and activate the chromatin to trigger genome activation. We show that histone acetylation is essential to activate the cascade of genes responsible to induce transient totipotency, trigger the program of maternal mRNA decay through miR-430, and initiate zygote development.

## Acknowledgments

We thank K. Bishop for technical help; K. Bilguvar, S. Mane and C. Castaldi for sequencing support; C. Kontur, V Tornini, and all the members of the Giraldez laboratory for critical reading of the manuscript, intellectual and technical support. JL Gomez-Skarmeta and JR Martinez-Morales for sharing ATAC seq before publication. David Gonzalez, Al Mennone and Joerg Bewersdorf for mentorship in image analysis. The imaging work was supported in part by NIH grant S10 OD020142. This research was supported by the Fonds de Recherche du Québec - Santé (postdoctoral fellowship to JDB), and the National Institute of Health (grants R01 HD074078, GM103789, GM102251, GM101108 and GM081602, R35 GM122580, 4DNucleome program to AJG). MAMM is supported by Programa de Movilidad en Areas de Investigación priorizadas por la Consejería de Igualdad, Salud y Políticas Sociales de la Junta de Andalucía. The Giraldez lab is supported by the Howard Hughes Medical Institute Faculty Scholar program, and was supported by the Pew Scholars Program in the Biomedical Sciences, March of Dimes 1-FY12-230, the Yale Scholars Program and Whitman fellowship funds provided by E. E. Just, Lucy B. Lemann, Evelyn and Melvin Spiegel, The H. Keffer Hartline and Edward F. MacNichol, Jr. of the Marine Biological Laboratory in Woods Hole, MA to AJG.

## Author Contributions

Author Contributions SHC and AJG conceived the project. SHC performed embryonic experiments and image analysis with the support of HDC and MAMM. YT performed RNA-seq analysis, with the contribution of CEV. LM, DM, performed Western blots and DNA measurements, SHC and JDB performed Click-iT-seq, SHC and HDC performed Click-iT staining, SHC and JPF performed chemical drug screening, and CEV performed data processing. SHC, YT, MAMM and AJG performed data analysis and, together with the other authors interpreted the results. AJG supervised the project, with the contribution of MAMM. SHC, MAMM and AJG wrote the manuscript with input from the other authors.

The authors declare no competing financial interests. Readers are welcome to comment on the online version of the paper. Correspondence and requests for materials should be addressed to AJG and MAMM.

## Methods

### Zebrafish maintenance and mating

Zebrafish wild-type embryos were obtained from natural matings of 12-month-old adult zebrafish of mixed wild-type backgrounds (TU-AB, and TL strains). Wild-type adults were selected randomly for mating. Zebrafish were maintained in accordance with AAALAC research guidelines, under a protocol approved by Yale University IACUC. All zebrafish and embryo experiments were carried out at 28°C.

### Constructs, sgRNA design, and mRNA and sgRNA in vitro transcription

Zebrafish *chk1* ORF was amplified from cDNA from 64 cell-stage embryos using primers 5’-TTTTCCATGGCTGTGCCTTTTGTTAAAG-3’ and 5’-TTTTCCGCGGTCAAATCAATGGCAAAACCTTTTGG-3’. The resultant PCR product was digested with restriction enzymes *NcoI* and *SacII* and ligated into the plasmid pT3TS-zCas9^1^. The final construct was confirmed by sequencing and corresponded in sequence to chk1 protein XP_021324451.1. pCS2-Smarca4a (zebrafish ORF) plasmid was a gift from the Cairns lab (University of Utah). dCas9-3xGFP was generated as follows: 3xGFP was PCR amplified from plasmid pHAGE-TO-dCas9-3XGFP2 using primers 5’ TTTCCGCGGCTACTCGAGTTTGTACAGTTC-3’ and 5’-TTTACCGGTGAGATCTCCTAAGAAGAAGAGAAAGGTGGGCTCTACTAGTGGCTCT-3’. pHAGE-TO-dCas9-3XGFP was a gift from Thoru Pederson (Addgene plasmid # 64107). PCR products were digested with restriction enzymes *AgeI* and *SacII* and ligated into the pT3TS-dCas9 plasmid, which is based on pT3TS-zCas9, containing point mutations that catalytically inactivate Cas9, also called dead Cas9 (dCas9)^3,4^. sgRNAs against the miR-430 locus were designed using an updated version of the CRISPRscan (crisprscan.org) tool^5^. Two different sgRNAs were used in combination with dCas9-3XGFP to label miR-430 locus: sgRNA 1 specific oligonucleotide: 5’-atttaggtgacactataGAGGGTACC-GATAGAGACAA<u>gttttagagctagaa</u> and sgRNA 2 specific oligonucleotide: 5’-taatacgactcactataGGCTGAGTGTTAACGACTG<u>gttttagagctagaa</u>. sgRNA 1 and sgRNA 2 target 11 and 9 sites in the miR-430 locus at chromosome 4, respectively. sgRNAs were generated as previously described^5,6^. Briefly, a 52-nt oligo containing the SP6 (5’-atttaggtgacactata) (sgRNA 1) or T7 (5′-taatacgactcactata) (sgRNA2) promoter, 20-nt of specific sgRNA DNA-binding sequence, and a constant 15-nt tail (small case and underlined) for annealing was used in combination with an 80-nt reverse universal oligo to add the sgRNA invariable 3′ end (5′-AAAAGCACCGACTCGGTGCCACTTTTTCAAGTTGATAACGGACTAGCCT-TATTTTAACTTGCTATTTCTAGCTCTAAAAC). A 117-bp PCR product was generated and purified using QIAquick PCR purification kit (Qiagen). The purified product was used as a template for T7 or SP6 *in vitro* transcription (AmpliScribe-T7-Flash transcription kit from Epicenter; MAXIscript SP6 Transcription Kit from Thermo Fisher Scientific, over a 6–7 h of reaction). *In vitro* transcribed sgRNAs were DNAse-treated, precipitated with sodium acetate/ethanol and checked for RNA integrity on a 2% agarose gel stained with ethidium bromide. To generate dCas9-3xGFP, Chk1, and Smarca4a capped mRNAs, the DNA templates were linearized using *XbaI* (dCas9-3xGFP and Chk1) or *SacII* (Smarca4a) and mRNA was synthetized using the mMessage mMachine T3 or SP6 (Smarca4a) kit (Ambion). *In vitro* transcribed mRNAs were DNAse I treated and purified using the RNeasy Mini Kit (Qiagen). Capped mRNA of Nanog, SoxB1, and Pou5f1 were prepared as previously described^7^.

### DNA quantification by qPCR

5 embryos were manually deyolked in Ringer’s solution (116mM NaCl; 1.8mM CaCl_2_, 2.9mM KCl; 5ml HEPES). Genomic DNA was extracted with PureLink Genomic DNA Kits (Thermo Fisher Scientific) following manufacture’s instruction and eluted with 500 μl of MilliQ water. To quantify nuclei, 3 μL of the genomic DNA was used in a 20 μL reaction containing 1 μL of primers amplifying repetitive miR-430 loci (Forward: 5’-CAAATGTGTGAAAAATCCCATC-3’; Reverse: 5’-AAGGGTGCACTTGCCTTATG-3’), using power SYBR Green PCR Master Mix Kit (Applied Biosystems) and a ViiA 7 instrument (Applied Biosystems). PCR cycling profile consisted of incubation at 50 °C for 2 min, followed by a denaturing step at 95 °C for 10 min and 40 cycles at 95 °C for 15 s and 60 °C for 1 min.

### Embryo injections and treatments

All injections and drug treatments were carried out on wild-type one-cell stage dechorionated embryos, unless otherwise noted. Experimental samples were then collected at the specified developmental stages/time as described in the text and figure legends. Varying amounts of mRNA per embryo were used as follows: 160 pg (Chk1), 25 pg (dCas9-3XGFP), 50 pg (Smarca4a, Nanog), 20 pg (SoxB1) and 30 pg (Pou5f1). P300 and BRD4 proteins were purchased from Protein One (P2004-01) and Reaction Biology Corp. (RD-21-153), respectively, and 200 pg of P300 and 630 pg of BRD4 protein were injected into each embryo. JQ1 treatment: embryos were bathed in 43.8 μM JQ1 (1:100 dilution from 4.38mM working stock in DMSO) to inhibit BET family of bromodomain proteins including BRD2, BRD3 and BRD4. SGC-CBP30 treatment: embryos were bathed in 20 μg/ml SGC-CBP30 (Sigma-aldrich, 1:500 dilution from 10mg/ml working stock in DMSO) to inhibit the bromodomain-containing transcription factors CREB binding protein (CBP) and Histone acetyltransferase p300 (EP300). Pol II inhibition: embryos were bathed in 5.8 μM triptolide (1:1,000 dilution from a 5.8 mM working stock in DMSO) or injected with 0.2ng of alpha-amanitin at one-cell stage to inhibit RNA polymerase II. Translation inhibition: embryos were collected at the one-cell stage. To limit the amount of translation of maternal mRNAs, embryos were transferred to media containing 10 μM pateamine A (PatA, purchased from D. Romoat Baylor University) and 50 μg/mL cycloheximide (CHX, Sigma Aldrich) at the 8-cell stage or the 32-cell stage, respectively. Because these treatments stop development, collection of these embryos was timed by monitoring the development of untreated sibling embryos when they reached the appropriate stage in this case sphere stage (4hpf). Chk1 treatment: 160pg of Chk1 mRNA was injected in one-cell stage embryos to inhibit/slow down DNA replication by inducing degradation of the limiting replication initiation factor Drf18. Embryos were collected when untreated sibling embryos reached the appropriate developmental stages: 64-cell stage (2hpf) or sphere stage (4hpf).

### dCas9-Labeling of endogenous miR-430 locus

To label the miR-430 locus, the two previously described sgRNAs were injected at 100pg each in combination with 25pg dCas9-3xGFP into dechorionated embryos at the one-cell stage. Embryos were image either live or fixed in 4% paraformaldehyde and processed for antibody staining against GFP.

### miR-430 transcription labeling by live imaging

To visualize nascent transcription of miR-430 by live imaging, a molecular beacon (MB^miR430^ : Dabcyl-5’-GCTGAACAGAGGTGACTAAGTCAGC-3’-Lissamine) was specifically designed to target the primary miR-430 transcript. The molecular beacon (MB^miR430^) was obtained from Gene Tools and resuspended in nuclease-free water. The molecular beacon was designed with a stem-loop structure, where the single-stranded loop region (5’-ACAGAGGTGACTAAG-3’) is antisense to the repetitive single stranded regions of the predicted structure of the primary miR-430 transcript. The single-stranded loop region is flanked by two 5-nt sequences (5’-GCTGA-3’ at the 5’ end and 5’-TCAGC-3’ at the 3’ of the molecular beacon) that are self-complementary and form the double-stranded stem. The molecular beacon is modified at the 5’ end with a fluorophore and the 3’ end with a quencher to improve the signal to noise ratio. 0.5pmole of MB^miR430^ was injected into wild type embryos at the one-cell stage. To trace individual nuclei during live imaging analysis, 0.4ng of Alexa Fluor^®^ 488 histone H1 conjugate (ThermoFisher Scientific, H-13188) was co-injected with MB^miR430^.

### Labeling zygotic transcription with uridine analogs for imaging

Click-iT RNA Alexa Fluor 594 Imaging Kit (C10330) was adapted for application in zebrafish embryos. Embryos were injected with 50 pmols of Click-iT^®^ RNA (5-ethynyl-uridine) (EU, E10345) or in kit form with Click-iT Nascent RNA capture kit, C-10365, and collected at the times/developmental stages indicated in the text and figure legends. After collection, embryos were fixed using a 4% paraformaldehyde (PFA) solution in 1x phosphate-buffered saline (PBS) at 4°C overnight. Fixed embryos were washed three times with 1x PBS and permeabilized using 0.5% Triton X-100 (PBS-T) at room temperature for a total of 30 minutes, followed by dehydration with serial dilutions of Methanol (25%, 50%, 75%, 100% Methanol diluted with 1x PBS-T). Dehydrated embryos were incubated at −20°C for at least 2 hours before rehydration with serial dilutions of Methanol (75%, 50%, 25%, 0% Methanol diluted with 1x PBS-T). Rehydrated embryos then underwent antibody staining, as described below, before proceeding with the manufacturer’s EdU-labeling visualization protocol. Briefly, embryos were incubated with a 1x working solution of Click-iT reaction cocktail, containing the Alexa Fluor 594 azide and CuSO_4_, for 1 hour in dark at room temperature. After removal of the reaction cocktail, embryos were washed once with Click-iT reaction rinse buffer. Embryos were then washed three times with 1x PBS-T and stained with DAPI, followed by dissection and mounting on glass slides in ProLong Diamond Antifade Mountant (ThermoFisher Scientific, P36965).

### Click-iT captured nascent zygotic transcripts for RNA-Sequencing

The Click-iT^®^ Nascent RNA Capture Kit (C10365) was adapted for application in zebrafish embryos. To capture nascent RNAs, 50 pmols of Click-iT^®^ EU (5-ethynyl Uridine) was injected in one-cell stage embryos and allowed to incorporate into the nascent zygotic transcript until the time point/developmental stage indicated in the text and figure legends. At these time points, total RNA from 35 embryos was extracted using TRIzol reagent (Invitrogen). The EU-incorporated RNAs were biotinylated and captured following the manufacturer’s instructions. Briefly, the EU-labeled RNA was biotinylated with 0.25mM biotin azide in Click-iT reaction buffer. The biotinylated RNAs were precipitated with etanol and resuspended in nuclease-free water. The biotinylated RNAs mixed with Dynabeads MyOne Streptavidin T1 magnetic beads in Click-iT RNA binding buffer and heated at 68°C for 5 minutes, followed by incubation at room temperature for 30 minutes while gently vortexing. The beads were immobilized using the DynaMag-2 magnet and were washed with Click-iT wash buffer1 and 2. The washed beads were then resuspended in Click-iT wash buffer2 and used for cDNA synthesis using the SuperScript^®^ VILO cDNA synthesis kit (Cat. no. 11754-050), followed with RNA-Sequencing. Single-stranded cDNA was recovered by heating the magnetic beads at 85°C for 5 minutes, ethanol precipitated, and dissolved in 20 μL of water. Libraries were constructed following Illumina TruSeq and dUTP protocol from cDNA. Sequencing was performed at the Yale Center for Genome Analysis and resulted into unstranded reads.

### Antibody staining and fluorescence microscopy

Embryos were fixed using a 4% paraformaldehyde (PFA) solution in 1x phosphate-buffered saline (PBS) at 4°C overnight. Fixed embryos were washed three times with 1x PBS and permeabilized using 0.5% Triton X-100 (PBS-T) at room temperature for a total of 30 minutes, followed by dehydration with serial dilutions of Methanol (25%, 50%, 75%, 100% Methanol diluted with 1x PBS-T). Dehydrated embryos were incubated at −20°C for at least 2 hours before rehydration with serial dilutions of Methanol (75%, 50%, 25%, 0% Methanol diluted with 1x PBS-T). Rehydrated embryos were washed with 1x PBS-T and incubated for 2-3 hours in blocking solution (1x PBS-T, 10% Bovine Serum Albumin), followed by overnight incubation at 4C with the primary antibodies specified in the text and figure legends. These primary antibodies include rabbit anti-Histone H3 (acetyl K27) antibody 1:1,000 (Abcam, ab177178), rabbit anti-RNA polymerase II CTD repeat YSPTSPS (phospho S5) antibody 1:1,000 (Abcam, ab5131), mouse anti-GFP Tag antibody 1:1,000 (ThermoFisher Scientific, A-11120). After three washes with 1x PBS-T, the embryos were incubated with secondary antibodies for 2 hours at room temperature. The secondary antibodies include Goat anti-Mouse IgG (H+L) secondary antibody, Alexa Fluor Plus 488 1:1,000 (ThermoFisher Scientific A32723), goat anti-rabbit-IgG-Atto647N 1:1,000 (Sigma-aldrich, 40839) and goat anti-mouse-IgG-Atto594 1:1,000 (Sigma-aldrich, 76085). All antibodies were diluted with blocking solution (1x PBS-T, 10% Bovine Serum Albumin). After this step, embryos were washed three times with 1x PBS-T before proceeding with either DAPI staining and mounting or treatment with Click-iT RNA Alexa Fluor 594imaging kit (ThermoFisher C10330) according to the manufacturer’s protocol. Samples were mounted in in ProLong Diamond Antifade Mountant (ThermoFisher Scientific, P36965) and examined using a confocal fluorescence microscope (Leica TCS SP8) and a Stimulated emission depletion (STED) microscope (Leica TCS SP8 Gated STED 3x super resolution microscope). Sequential imaging is used to avoid spectral bleedthrough in experiments involving multi-fluorescent staining. Fluorescence minus one (FMO) control is performed to configure image acquisition setting. In Click-iT labeled imaging analysis, Triptolide or α-amanitin treated samples will be used as a negative control for optimizing acquisition setting. DAPI staining is performed on all imaging samples for proper staging for embryos with comparable nuclei stages (Extended data figure 2b). Embryos with comparable nuclei stages within the same developmental stages are imaged and compared accordingly. The reported number of imaged nuclei comes from a minimal of three independent embryos. Live embryos are mounted in 0.4% low melt agarose (AmericanBio, CAS: 9012-36-6) and all live imaging is performed at monitored temperature of 28°C. Images were processed and quantified with Bit-plane Imaris, Image-J software. Figure panels display maximum projections, unless specified as a single focal plane image for better presentation of co-localization across different channels.

### Western blot

Ten embryos (H3, H2B n=5; H2A, H4) were manually deyolked, snap frozen in liquid nitrogen and boiled at 95 °C for 5 min in 15 μL of water, 7.5 μL of 4x NuPAGE LDS Sample Buffer (ThermoFisher Scientific), 3 μL DTT (Sigma-Aldrich). Samples were run on 4–12% polyacrylamide NuPAGE Bis-Tris gels (ThermoFisher Scientific) for 45 min at 180 V and wet electrotransferred onto a nitrocellulose membrane (GE LifeSciences) for 70 min at 30 V. Membranes were incubated in blocking solution (5% milk in PBS-T) for 2 h. Then primary antibodies were diluted in blocking solution and incubated with the membrane overnight at 4°C Anti-H3 1:10,000 (ab1791; Abcam), H4 1:1,000 (ab10158; Abcam), H2A 1:1,000 (ab18255; Abcam), H2B 1:3,000 (ab1790; Abcam), Actin 1:5,000 (ab8227, Abcam). Secondary antibody Goat Anti-Rabbit IgG Antibody, (H+L) HRP conjugate (AP307P; Millipore) was incubated with the membrane at 1:10,000 for 1 h at RT. Membranes were analyzed by chemiluminescent detection and X-ray film (E3012, Denville Scientific).

### RNA-seq analysis and normalization

Total RNA from 20 embryos per condition at indicated time point/developmental stage were snap frozen in Liquid Nitrogen and the RNA was extracted using Trizol reagent (Invitrogen). Samples were treated with Epicentre Ribo-Zero Gold, to deplete ribosomal RNA, or subject to pull-down by oligo dT beads, to enrich for poly(A)+ RNA. TruSeq Illumina RNA sequencing libraries were constructed and samples were multiplexed and sequenced on Illumina HiSeq (High output) machines to produce 75-nt single-end reads by the Yale Center for Genome Analysis. Raw reads were mapped to zebrafish GRCz10 genome sequence using STAR v2.5.3^9^ with parameters *–alignEndsType Local* and *–sjdbScore 2*. Genomic sequence index for STAR was built including exon-junction coordinates from Ensembl v90^10^. Read counts per gene were computed by summing the total number of reads overlapping at least 10 nucleotides of the gene annotation. Gene models were constructed by merging all overlapping transcript isoforms of each gene. For miR-430, reads overlapping the locus on chromosome 4 from coordinate 28,738,727 to 28,754,891 were counted as miR-430 cluster reads. All reads mapping to all zebrafish genes, including reads mapping to multiple loci in the zebrafish genome, were kept. To compute read counts per gene, each locus where a read was mapped was assigned a weight equal to 1 divided by the total number of loci to which the read was mapped to. RNA reads were normalized to the total number of reads mapped to the zebrafish genome per kilobase per million. For experiments involving Click-iT-seq reads were normalized by mitochondrial reads mapping to the mitochondrial protein coding genes. Mitochondrial RNAs provides a valuable internal control as shown by the Neugebauer lab^11^, because i) it continues to be transcribed during the MZT^11^, ii) these RNAs are easily labeled with Click-iT, and iii)they can be used as an internal control because the mitochondrial RNA Pol II is not inhibited by the concentrations of -amanitin used in this study^12-14^.

### Determination of zygotic and maternal-zygotic genes

To calculate per gene RPKMs, the number of reads mapped to each gene from the Click-iT RNA-seq experiment were summed and normalized by gene length and the total number of reads mapped to the mitochondrial protein-coding genes, unless otherwise specified. Intron regions were defined as genic regions that are not covered by any extended exon (exon extending 15nt on both ends) on the same or any other gene. Only uniquely mapped reads were used for intron analysis. All zygotic and maternal zygotic genes were defined using criteria in the following table. Genes in each category must satisfy all the criteria in the column.

The criteria are detailed as follows (Extended Data Fig. 10):

- *Zygotically expressed genes from previous studies.* The union of all identified zygotically expressed zebrafish genes reported by the Neugebauer lab^11^ and the Giraldez lab^7^.
- *WT 4h Click-iT compared to WT 2h mRNA-seq R0.* This comparison was designed to distinguish between zygotic and maternal zygotic genes. Zygotic genes are expected to be enriched by Click-iT RNA-seq at 4 hpf compared to RiboZero purified maternal mRNAs at 2 hpf measured by mRNA-seq. In this comparison, read counts from the Click-iT RNA-seq experiment were normalized to RPKMs using the total exonic reads in the sample (figure S1b, e).
- *Exon RPKM in WT 4h Click-iT (normalized by total).* To define additional high confidence zygotic and maternal zygotic genes, a high Click-iT RPKM cut-off was applied. This is aimed to avoid potential background from maternal mRNAs purified by Click-iT. Cut-off of 10 RPKM was determined by minimum expression of zygotic genes identified in previous studies (figure S1a).
- *WT 4h Click-iT compared to triptolide 4h Click-iT.* Zygotic and maternal zygotic genes are expected to have higher exon and intron RPKM in WT embryos compared to triptolide treated embryos (figure S1b,c,e,f). Intron signal is more sensitive than exon signal as most maternally deposited mRNAs are spliced and the introns are degraded early on^7^. In this comparison, exonic and intronic read counts were normalized to RPKMs using the total reads mapping to mitochondrial protein-coding genes. As the overall intron RPKMs are ~4-10 fold lower than exon RPKMs (figure S1f), a 30 RPKM intron cut-off was used corresponding to ~7-fold decrease to the minimum RPKM cutoff used for exons.
- *WT 4h compared to triptolide 4h in mRNA-seq R0 on exon.* The additional zygotic and maternal zygotic candidates were further restricted by mRNA-seq as zygotically expressed genes were expected to have greater exon expression in WT embryos compared to embryos treated with triptolide.
- *WT 4h compared to triptolide 4h in mRNA-seq R0 on intron.* Zygotically expressed genes were expected to have greater intron expression as newly transcribed zygotically expressed genes are spliced and detectable in WT embryos, unless the gene has no intron or the intron is small, hence this condition is only applied if the intron length is >= 500bp.

### Identification of zygotically activated genes

For Click-iT RNA-sequencing assays in which ZGA factors were overexpressed (P300, BRD4), a gene was considered to be activated in the overexpression condition if the gene had at least 4-fold increase in exon or intron RPKM in the overexpression condition relative to triptolide or -amanitin treated embryos at the same stage. For Click-iT RNA-sequencing assays involving chemical or genetic treatment (chk1, Pateamine A+ Cycloheximide), genes were considered to be activated in the wildtype or treated embryos if those genes had at least 4-fold increase in exon RPKM or intron in the condition relative to the same condition treated with triptolide or -amanitin. The fold increase was calculated after adding 0.1 RPKM to genes in both conditions in the comparison for a more stringent filter for genes with low RPKM, which effectively sets a 0.3 RPKM cut off. For biplots created in this study, a small 0.1 RPKM was added to the expression of each gene in both conditions to allow the display of genes with zero counts in any condition. For biplots comparing gene expression in chk1 or PatA+CHX treated embryos, 0.01 RPKM was added to each gene given that these embryos have lower gene expression due to the fewer number cells and DNA template per embryo.

### ChIP-seq analysis of histone marks

Previously published ChIP-seq data [GSE32483] of H3K27ac, H3K4me3 and H3K4me1 marks at dome stage were realigned to the zebrafish GRCz10 genome using Bowtie v1.1.2^15^ using parameters -v 3 –best –strata –all –chunkmbs 1,000 -m 2,000. Histone mark signal for each gene was calculated by the total number of reads mapped to the gene body region and 1,000 nt upstream of the transcription start site, and normalized by the total number of reads aligned the genome and the length of the gene body plus 1,000 nt upstream. Histone mark signal per Kilobase (RPKM) for each gene was reported and used for the correlation with gene transcription level. The gene length for miR-430 is 17164nt = 16164 + 1000nt.

### Heatmap

Heatmaps were created using R 3.3.3 and package gplots. Fold change of gene expression between different conditions was first taken base 2 logarithm and then capped at −5 to 5. Genes with no expression in one of the conditions in the comparison were assigned infinite fold change and capped by the value mentioned above. Genes with no expression in both of the conditions were taken as no change between conditions.

---

**Extended data Figure 1.**
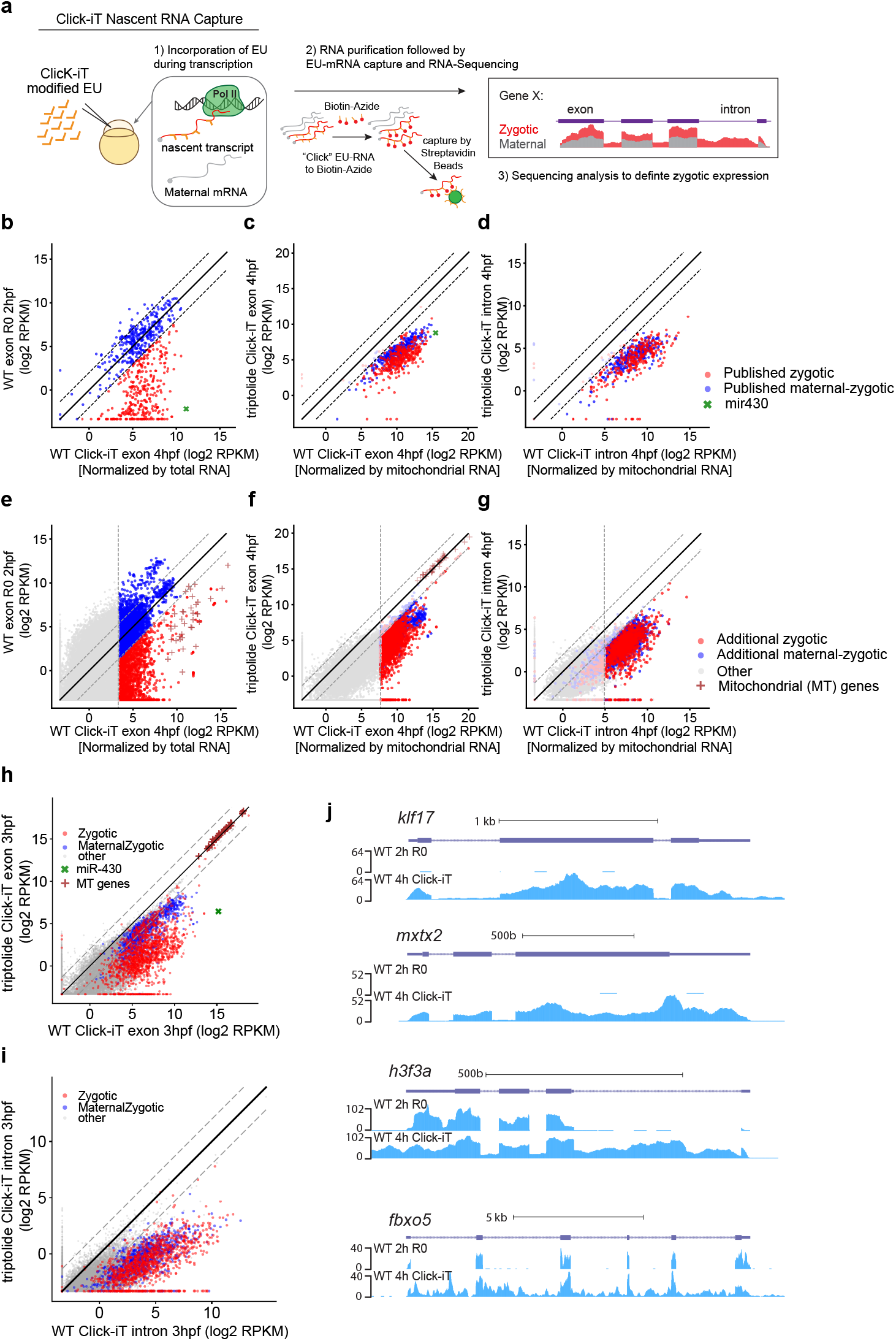
Identifying de novo zygotic transcription by Click-iT-seq. **(a)** Schematic illustrating the workflow of Click-iT pulldown assay to capture nascent RNA during ZGA for RNA-sequencing (Click-iT-seq). **(b)** Biplot comparing exon expression levels of zygotic and maternal-zygotic genes defined in published studies (Methods) from Click-iT-seq experiment at 4hpf and mRNA-seq experiment at 2hpf in wild-type embryos. **(c-d)** Biplot comparing expression levels of exons (c) and introns (d) of zygotic and maternal-zygotic genes defined in published studies (Methods) in wild-type (WT) and triptolide treated embryos (triptolide) from Click-iT-seq experiment at 4hpf. **(e)** Biplot comparing exon expression levels of additional zygotic and maternal-zygotic genes (Methods) between Click-iT-seq experiment at 4hpf and mRNA-seq experiment at 2hpf in wild-type embryos. **(f-g)** Biplot comparing expression levels of exons (f) and introns (g) of additional zygotic and maternal-zygotic genes (Methods) between wild-type (WT) and triptolide treated embryos (triptolide) from Click-iT-seq experiment at 4hpf. **(h-i)** Biplot comparing expression levels of exons (h) and introns (i) of genes between wild-type (WT) and triptolide treated embryos (triptolide) from Click-iT-seq experiment at 3hpf. **(j)** Genome tracks representing normalized mRNA-seq and Click-iT-seq signal measured at 2hpf and 4hpf in wild-type embryos for 2 examples of zygotic genes (*klf17* and *mxtx2*) and 2 examples of maternal-zygotic genes (*h3f3a* and *fbxo5*). In (b-i), dashed lines represent 4-fold change. Here, 4-fold change in (b) and (e) serves as a criteria separating zygotic and maternal-zygotic genes, and 4-fold change in (c), (d), (f), and (g) serves as criteria for defining zygotically expressed genes (Methods). In (b) and (e), the RPKM from the Click-iT-seq experiment is derived from total read number mapped to the exons on the gene normalized by the total exonic reads in the same sample. In (c), (d), (f), and (g), the RPKM from the Click-iT-seq experiment is derived from total number of reads mapped to the gene normalized by the total number of reads mapped to mitochondrial protein-coding genes in the same sample.

**Extended data Figure 2.**
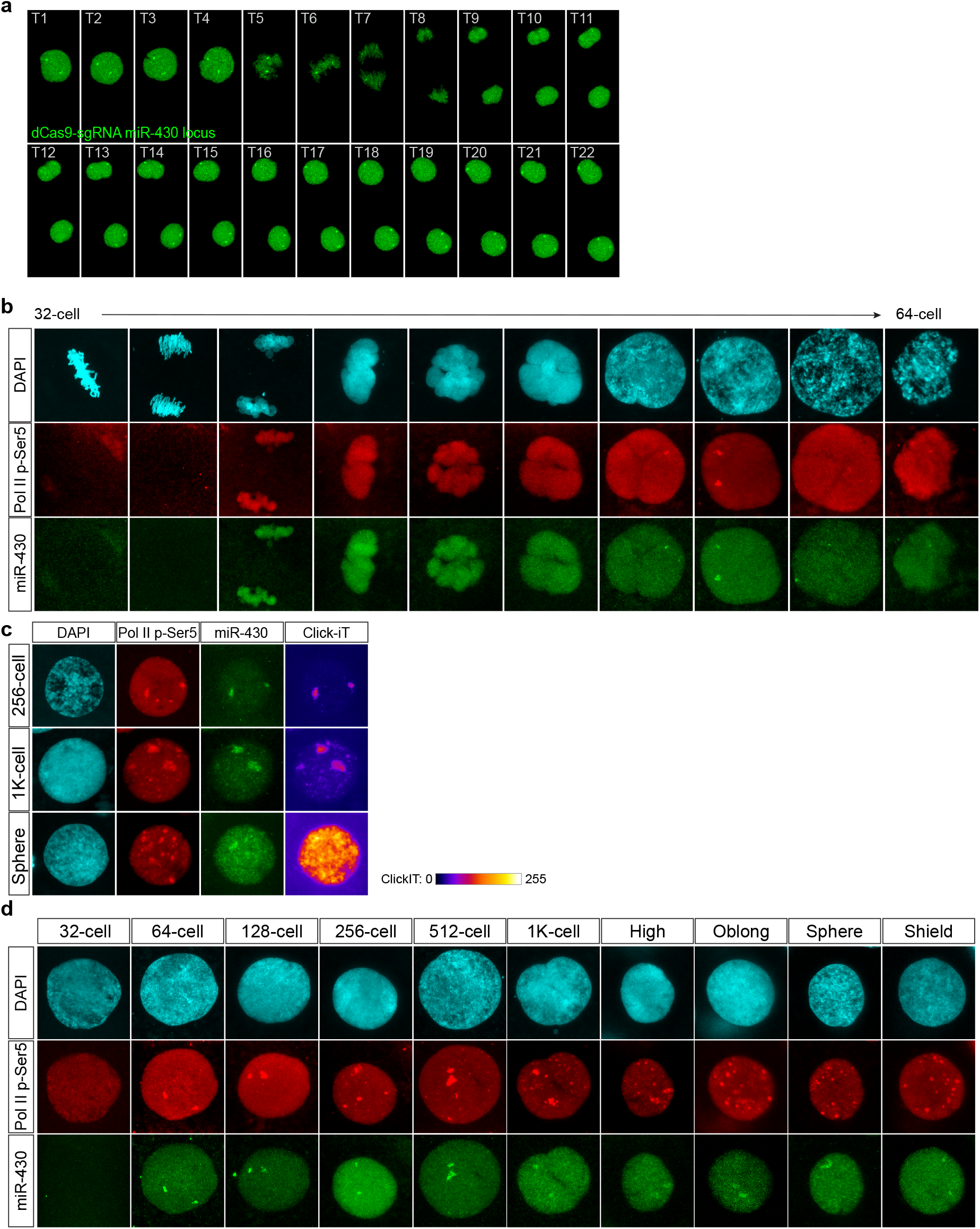
Temporal analysis of zygotic transcription by single nucleus imaging. **(a)** CRISPR-dCas9 labeled miR-430 locus in live zebrafish embryos over cell cycle. The temporal interval between each confocal image (T1-T22) is 1 minute. A mask is applied to the maximal projected confocal images to trace the displayed nuclei over time with minimal background. **(b)** Time-resolved confocal images labeled for DAPI, Pol II p-Ser5, miR-430 loci. Displayed nuclei are collected from embryos between 32-and 64-cell stages that are fixed in 2 minutes intervals throughout the cell cycle. Embryos with different temporal stages within the cell cycle was organized in a presumptive chronological order based on DAPI staining to illustrate the subset of nuclei stage when Pol II p-Ser5 can be detected throughout the rapid cell cycle. Note that Pol II p-Ser5 signal emerges as bright foci in late interphase and early prophase. **(c)** Representative single nucleus confocal images labeled with DAPI, Pol II p-Ser5, miR-430 loci and Click-iT at 256-, 1K-cell and Sphere stage. Note the co-localization of miR-430 loci with Pol II p-Ser5 signal as well as Click-iT signal. Click-iT signal intensity is presented in a heatmap color scale**. (d)** Time-course analysis of single nuclei labeled for Pol II p-Ser5 and dCas9 targeting the miR-430 locus from 32-cells to shield stage. Note that the first detectable Pol II p-Ser5 foci signal co-localizes with miR-430 loci at 64-cell stage. As transcription is globally activated, multiple Pol II p-Ser5 foci appear across the nucleus. DAPI staining is used to select temporally comparable stages of nucleus within each cell cycle as rationalized above.

**Extended data Figure 3.**
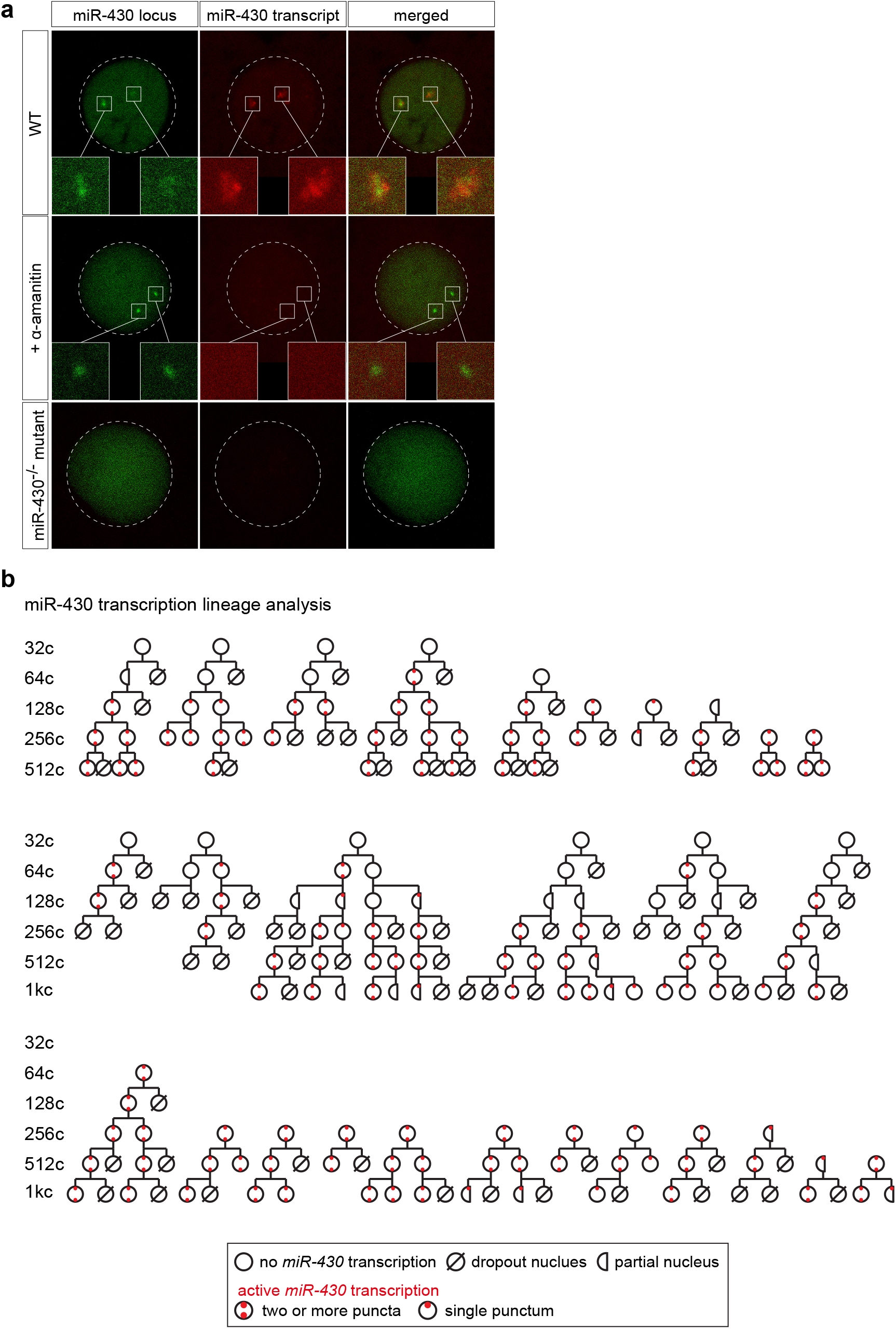
Cell-lineage tracing for miR-430 transcription with molecular beacon by live imaging. **(a)** Confocal imaging of wild-type (WT), α-amanitin treated (+α-amanitin), and miR-430^−/-^ mutant embryos labeled with MB^miR430^ (red) and dCas9-miR-430 (green). Note the absence of miR-430 transcription in miR-430^−/-^ deletion mutants and after Pol II inhibition. Also note the co-localization of MB^miR430^ signal with dCas9-miR-430 signal in wild-type embryos from the merged channel confocal image. **(b)** Cell-lineage tracing for miR-430 transcription. Nuclei from two independent developing live zebrafish embryos are imaged. Transcription competency is acquired in a stochastic manner at 64c, and once gained is maintained in the daughter cells after cell division. Transcription of miR-430 is scored with the detection of puncta signal from the molecular beacon (MB^miR430^). Nuclei dropped out from the field of view during live imaging were noted as either partially visualized (partial nucleus) or untraceable (dropout nucleus).

**Extended data Figure 4.**
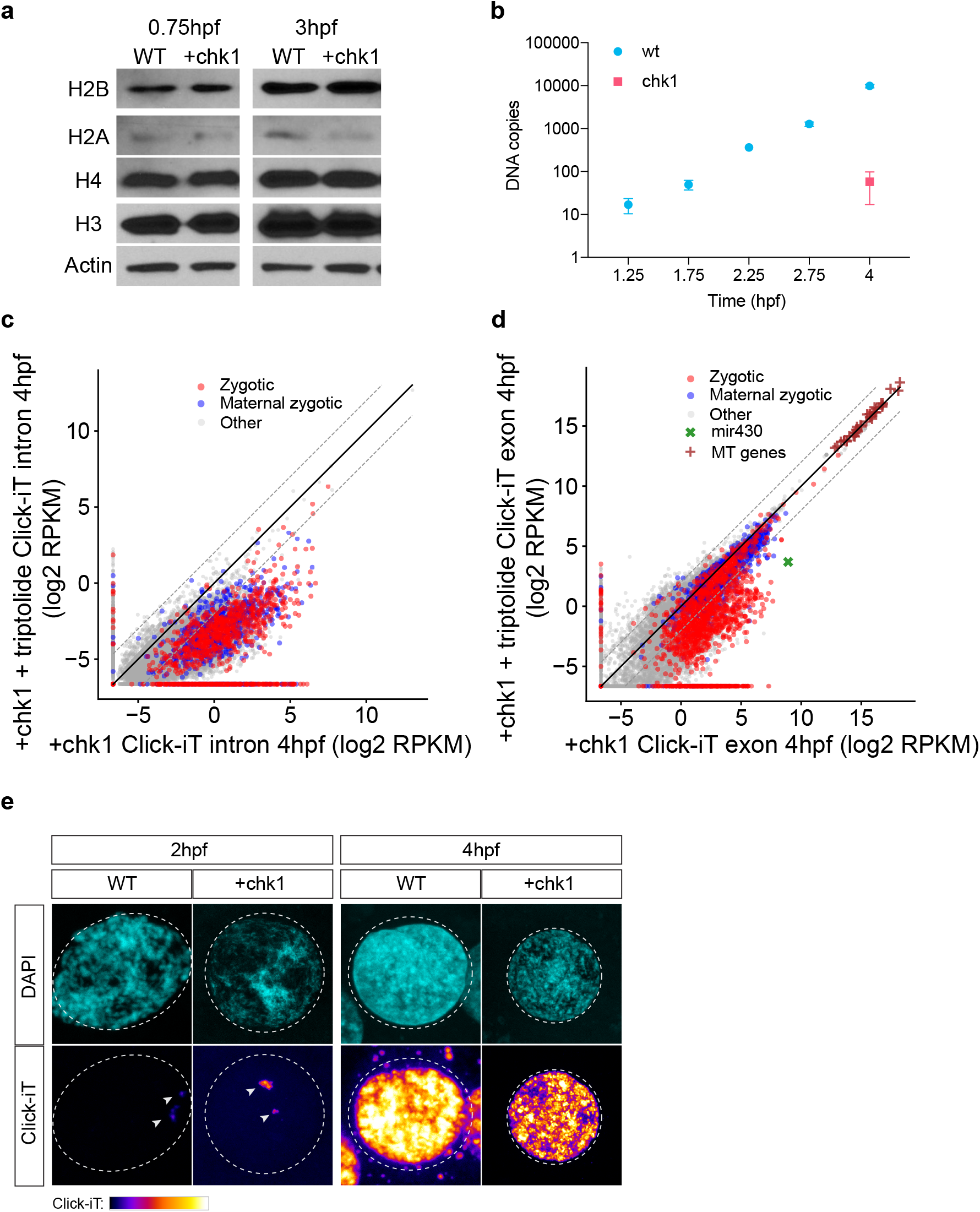
Zygotic genome activation occurs in chk1-injected embryos despite their lower N/C ratio. **(a)** Western blot of lysate from wild-type (WT) and chk1-injected (+chk1) zebrafish embryos collected at 0.75hpf and 3hpf respectively. Note the comparable histone (H2B, H2A, H3, H4) protein levels between the time-matched wild-type and chk1-injected embryos. Beta-Actin was used as loading control. **(b)** Quantification of DNA copies by qPCR analysis of miR-430 locus in 8-cell (1.25hpf), 32-cell (1.75hpf), 128-cell (2.25hpf), 512-cell (2.75hpf) and sphere stage (4hpf) wild-type embryos v.s. chk1-injected embryos collected at 4hpf. Note that the number of DNA templates for miR-430 in chk1-injected embryos at 4 hpf resembles that from embryos between 8-and 32-cell stages. 4 replicates were analyzed. **(c-d)** Biplot comparing intron (c) and exon (d) expression levels of genes in chk1-injected embryos with (+chk1+triptolide) and without triptolide (+chk1). Note that zygotic genome activation occurs in chk1-injected embryos at 4hpf. Dashed lines represent 4-fold change. **(e)** Single nucleus confocal image of DAPI and Click-iT in wild-type (WT) and chk1-injected (+chk1) embryo at 2hpf and 4hpf. Click-iT signals are highlighted by white arrows in the confocal images for 2hpf embryos. Click-iT signal intensity is presented in a heatmap color scale (2hpf: 0-255; 4hpf: 20-255). Note the increase in Click-iT signal intensity in chk1-injected (+chk1) embryos than wild-type (WT) embryos at 2hpf. Genome activation takes place in a time-dependent manner at 4hpf despite the lengthening of cell cycle during early development.

**Extended data Figure 5.**
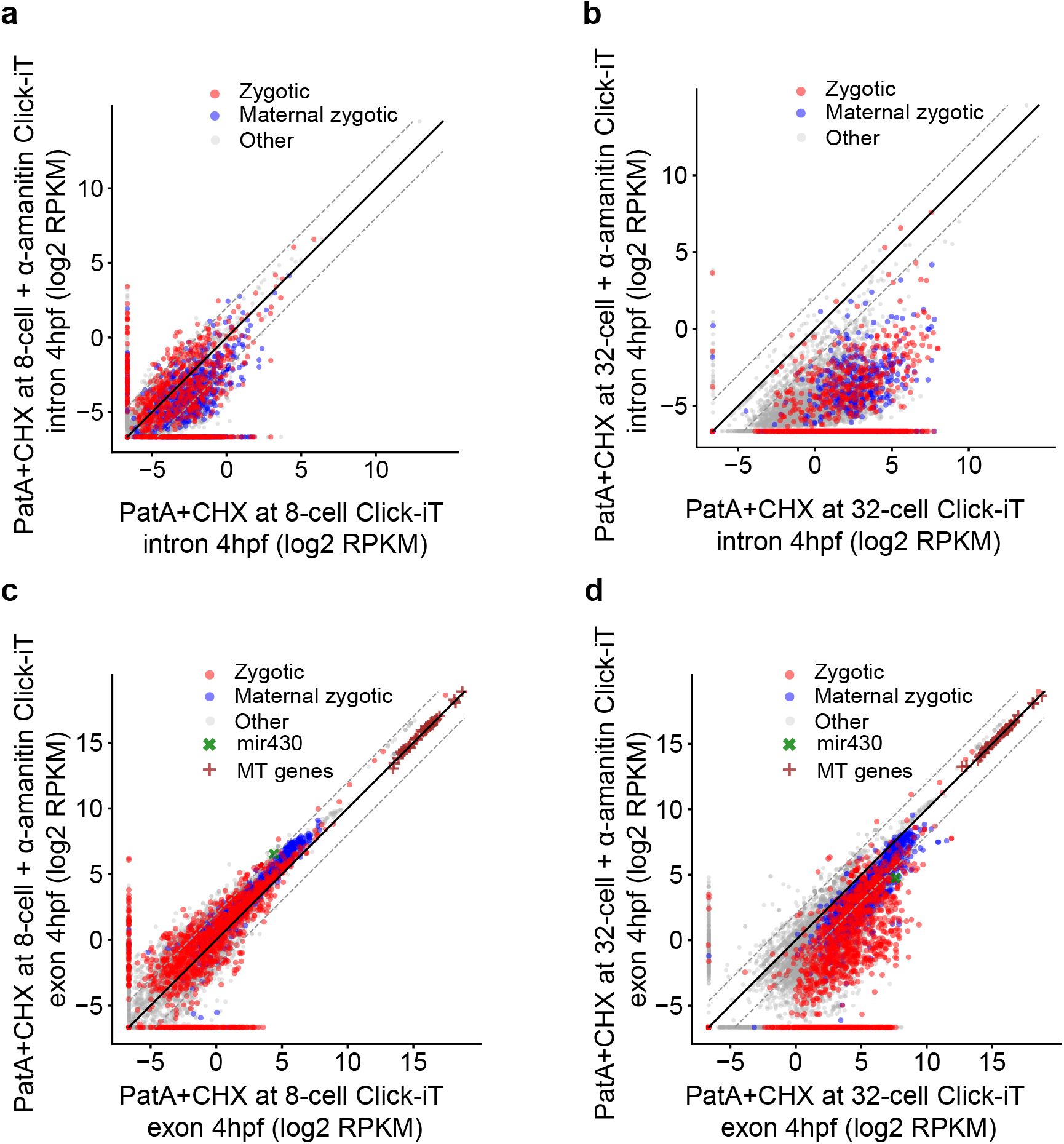
Embryos treated with PatA and CHX at 8-and 32-cell stages demonstrate different transcriptional competency at 4hpf. **(a-b)** Biplot of Click-iT-seq RNA levels at 4hpf comparing the level of transcription using intron signal. Embryos were treated with PatA + CHX at 8-cell stage (a) or at 32-cell stage (b), with and without α-amanitin. **(c-d)** Biplot of Click-iT-seq RNA levels at 4hpf comparing the level of transcription using exon signal. Embryos were treated with PatA + CHX at 8-cell stage (c) or at 32-cell stage (d), with and without α-amanitin. Note that embryos with PatA and CHX treatment at 32-cell stage demonstrate higher level of transcription competency than embryos with PatA and CHX treatment at 8-cell stage, indicating the additional time for translation of maternal RNAs is required for zygotic genome activation. Dashed lines represent 4-fold change. Intron expression comparison allows more sensitive measurement of zygotic RNA expression from maternal RNA contribution.

**Extended data Figure 6.**
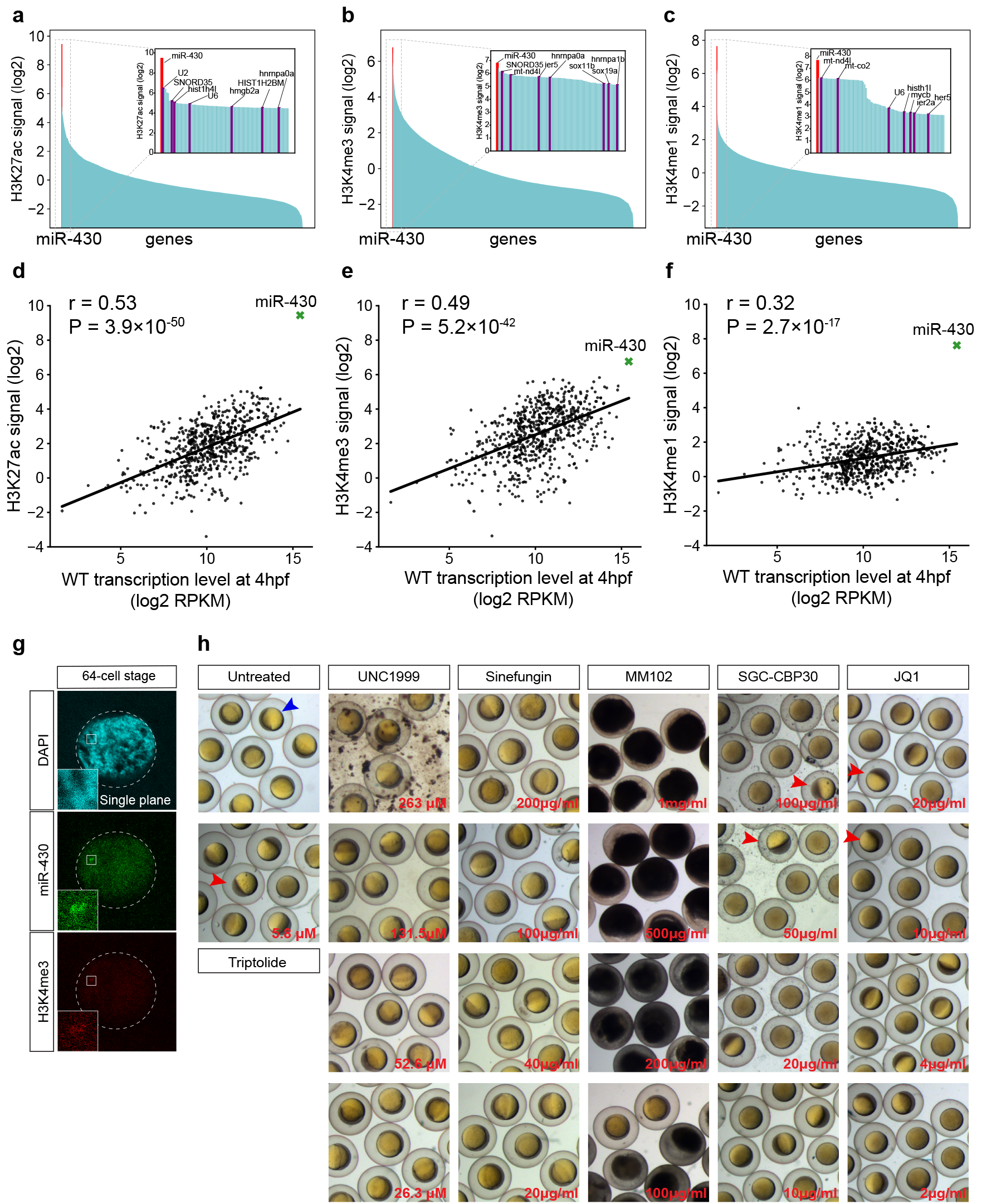
miR-430 is among the top labeled loci with H3K27ac, H3K4me3, and H3K4me1 marks. **(a-c)** Barplot showing level of H3K27ac (a), H3K4me3 (b) and H3K4me1 (c) on all zebrafish genes. Inset barplot represents the top-100 genes. MiR-430 locus is represented by a red bar. Note miR-430 is among the top labeled loci with H3K27ac, H3K4me3 and H3K4me1. **(d)** Biplot showing significant correlation between levels of H3K27ac (r = 0.53 (Spearman correlation), *P* = 3.9×10^−50^, rank correlation independence test) and gene transcription level (difference in gene expression (exon RPKM) between wild-type embryos and triptolide treated embryos) on published zygotic and maternal-zygotic genes (Methods). **(e)** Biplot showing significant correlation between levels of H3K4me3 (r = 0.49 (Spearman correlation), *P* = 5.2×10^−42^, rank correlation independence test) and gene transcription level. **(f)** Biplot showing significant correlation between levels of H3K4me1 (r = 0.32 (Spearman correlation), *P* = 2.7×10^−17^, rank correlation independence test) and gene transcription level. In (d-f), trend line is shown in the scatter plots. **(g)** Single plane stimulated emission depletion microscopy (STED) image labeled for H3K4me3, dCas9-miR-430 and DAPI (confocal). Note the co-localization of H3K4me3 with individual miR-430 locus. **(h)** The effect of titrated treatments of different small molecule inhibitors targeting readers or writers for H3K4me3/H3K4me1 and H3K27Ac on gastrulation at 6hpf. UNC1999 (Sigma Aldrich) is a selective inhibitor of both EZH2 and EZH1 lysine methyltransferases; Sinefungin (Abcam) is a competitive methyltransferase inhibitor; MM-102 (Selleck Chem.) is a selective WDR5/MLL (WD-repeat protein 5/mixed-lineage leukemia methyltransferase complex) interaction inhibitor; SGC-CBP30 (SGC) is a potent inhibitor for CREBP (cAMP-responsive element-binding protein binding protein) and EP300 (E1A-associated protein p300) bromodomains; JQ1 (Sigma Aldrich) is a selective bromodomain inhibitor targeting the BET (bromodomain and extra terminal domain) family of proteins, including BRD2, BRD3 and BRD4. Blue arrow denotes a gastrulating wild-type embryo at 6hpf; red arrows denote embryos with gastrulation arrest phenotype at 6hpf. Note that embryos with JQ1 and SGC-CBP30 treatment fail to gastrulate similar to the phenotype shown in triptolide treated embryos (Triptolide).

**Extended data Figure 7.**
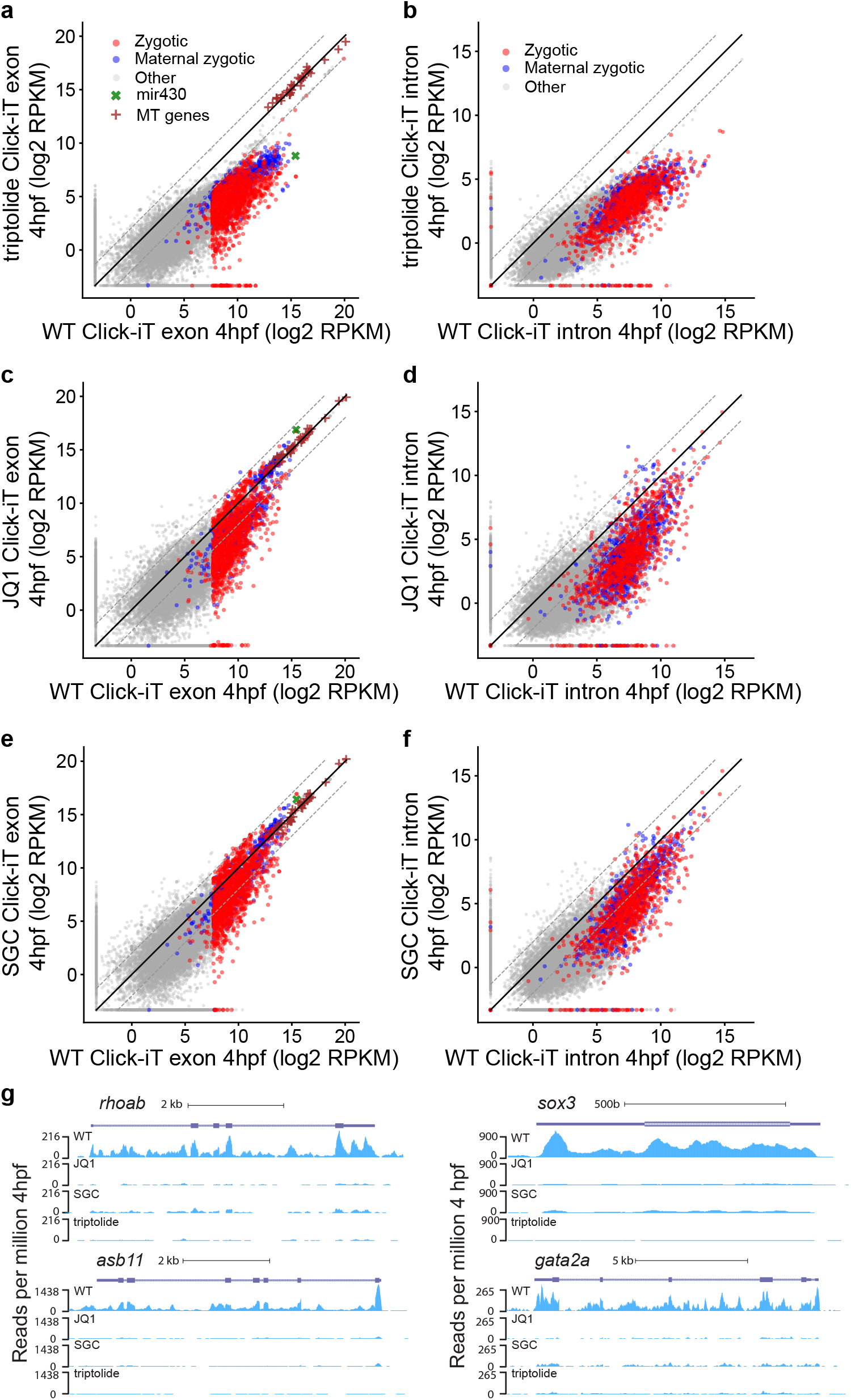
JQ1 and SGC treatment perturbs zygotic gene activation similar to triptolide treatment. **(a-b)** Biplot comparing exon (a) and intron (b) expression levels of genes measured by Click-iT-seq in triptolide treated embryos (triptolide) with wild-type embryos (WT) at 4hpf. Dashed lines represent 4-fold change. **(c-d)** Biplot comparing exon (c) and intron (d) expression levels of genes measured by Click-iT-seq in JQ1 treated embryos (JQ1) with wild-type embryos (WT) at 4hpf. **(e-f)** Biplot comparing exon (e) and intron (f) expression levels of genes measured by Click-iT-seq in SGC treated embryos (SGC) with wild-type embryos (WT) at 4hpf. **(g)** Genome tracks representing normalized Click-iT-seq signal measured at 4hpf in wild-type, triptolide, JQ1 and SGC treated embryos for examples of house keeping (*rhoab, asb11*) and developmental (*sox3, gata2a*) genes. Note the marked reduction in gene expression in both JQ1 and SGC treated embryos respectively when compared to wild-type embryos.

**Extended data Figure 8.**
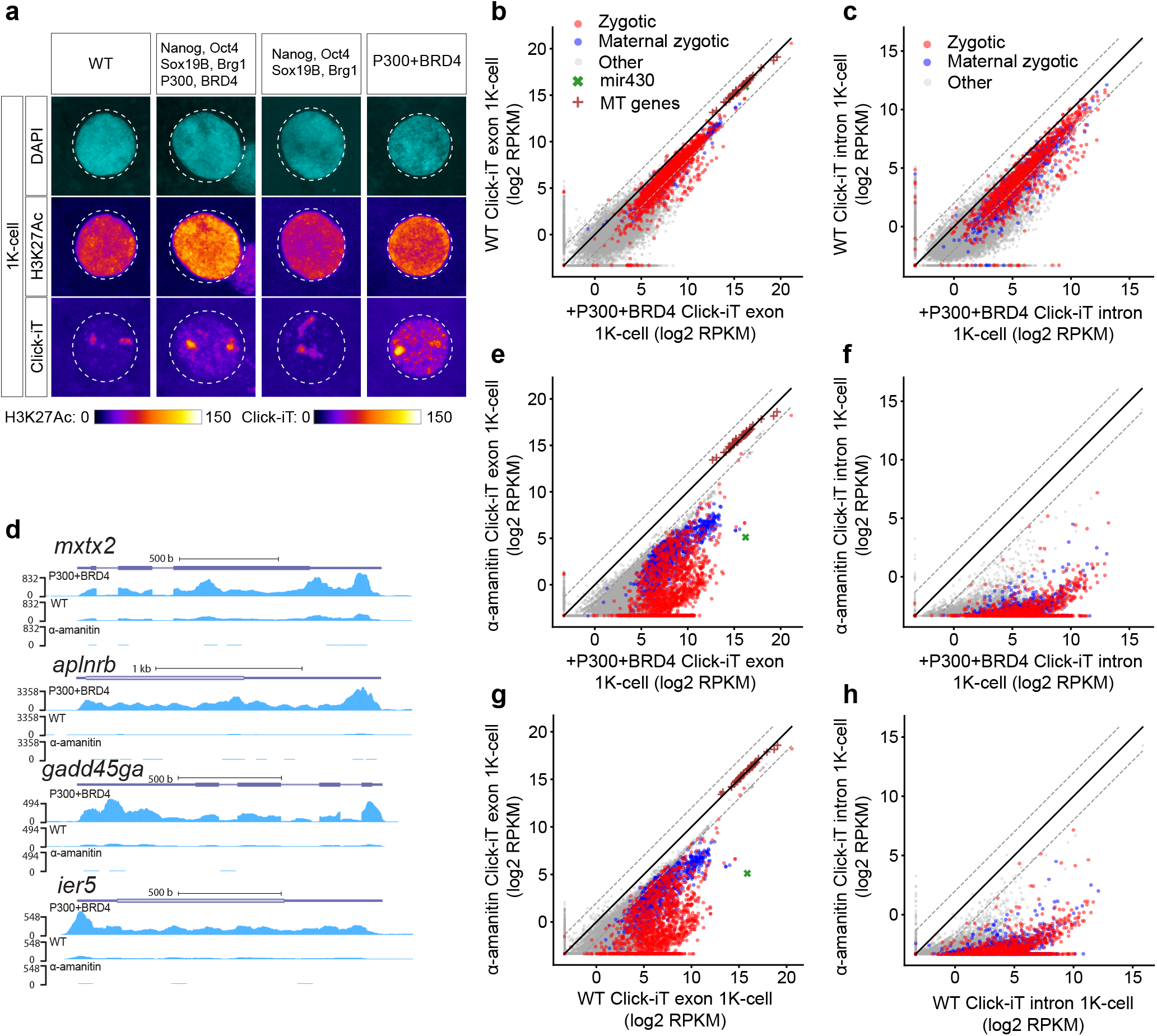
Early expression of P300 and BRD4 proteins leads to premature activation of the zygotic genome. **(a)** Single nucleus imaging analysis of H3K27Ac and Click-iT signal of embryos in different conditions at 1K-cell stage (WT: wild-type embryos; Nanog, Oct4, Sox19b, Brg1, P300, BRD4: embryos with early expression of Nanog, Oct4, Sox19b, Brg1, P300 and BRD4 proteins; Nanog, Oct4, Sox19b, Brg1: embryos with early expression of Nanog, Oct4, Sox19b, and Brg1 proteins; P300+BRD4: embryos with early expression of P300 and BRD4 proteins). Both H3K27Ac and Click-iT signal intensity are presented in a heatmap color scale. Note the increased H3K27Ac and Click-iT signal in embryos with early expression of Nanog, Oct4, Sox19b, Brg1, P300 and BRD4 proteins and embryos with early expression of P300 and BRD4 proteins when compared to wild-type (WT) embryos. **(b-c)** Biplot comparing exon (b) and intron (c) expression levels of genes measured by Click-iT-seq in wild-type (WT) embryos and embryos with early expression of P300 and BRD4 proteins (+P300+BRD4) at 1K-cell stage. Dashed lines represent 4-fold change. **(d)** Genome tracks representing normalized Click-iT-seq signal measured at 1K-cell stage in embryos with early expression of P300 and BRD4 (P300+BRD4), wild-type (WT) and α-amanitin treated (α-amanitin) embryos for examples of zygotic genes. **(e-f)** Biplot comparing exon (e) and intron (f) expression levels of genes measured by Click-iT-seq in embryos with early expression of P300 and BRD4 proteins (+P300+BRD4) and α-amanitin treated embryos (α-amanitin) at 1K-cell stage. **(g-h)** Biplot comparing exon (g) and intron (h) expression levels of genes measured by Click-iT-seq in wild-type embryos (WT) and α-amanitin treated embryos (α-amanitin) at 1K-cell stage. Note the marked increase in gene expression in embryos with early expression of P300 and BRD4 proteins when compared to wild-type embryos (WT).

**Extended data Figure 9.**
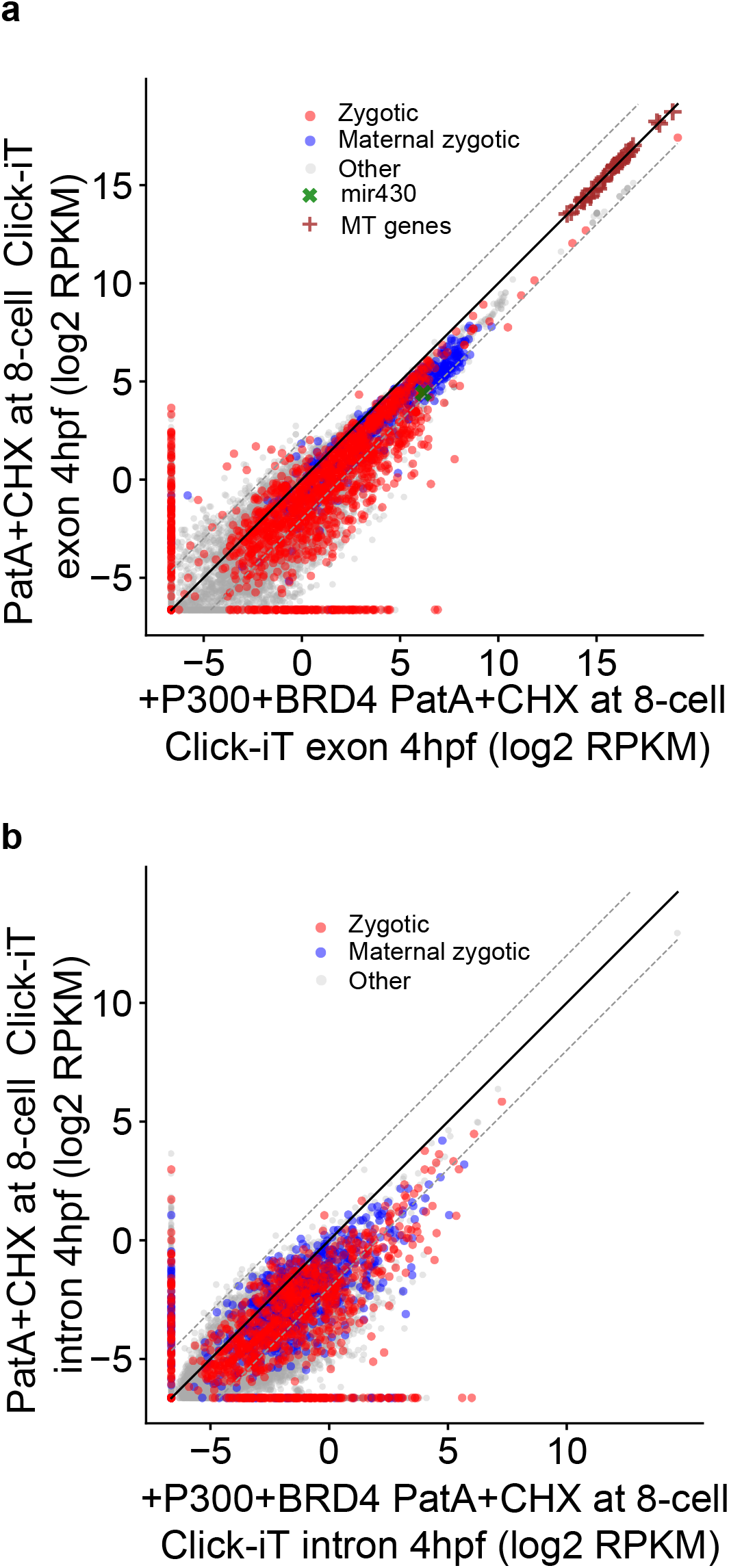
Overexpression of P300 and BRD4 protein restore transcription competency in embryos treated with PatA and CHX at 8-cell stage. **(a-b)** Biplot comparing exon **(a)** and intron **(b)** expression levels of genes measured by Click-iT-seq in embryos treated with PatA and CHX at 8-cell stage with (+P300+BRD4 PatA+CHX at 8-cell) and without early expression of P300 and BRD4 (PatA+CHX at 8-cell) at 4hpf. Dashed lines represent 4-fold change. Note the marked increase in gene expression in embryos treated with PatA and CHX at 8-cell stage with early expression of P300 and BRD4 when compared to embryos treated with PatA and CHX at 8-cell stage at 4hpf.

**Extended data Figure 10.**
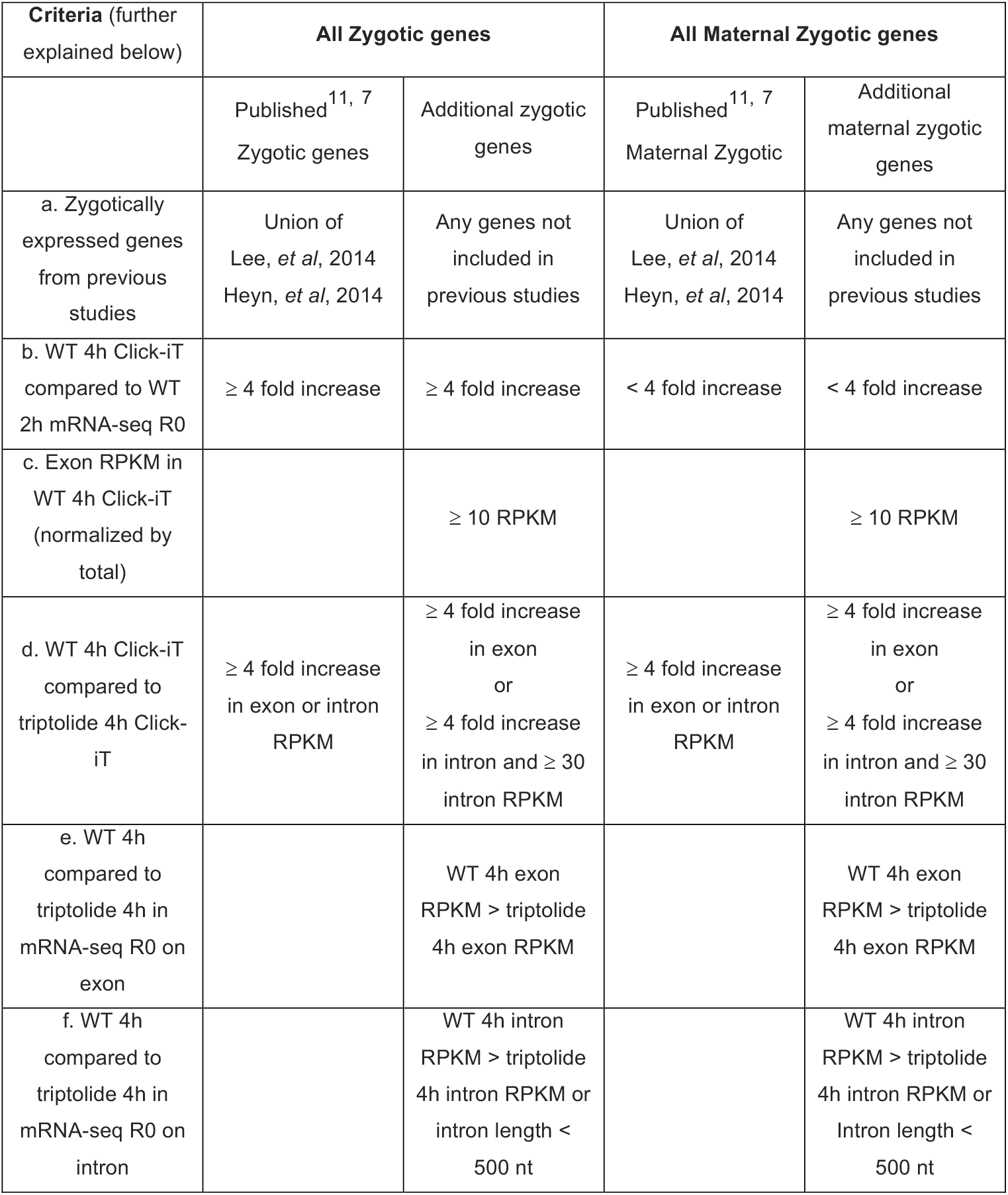
Determination of zygotic and maternal-zygotic genes.

